# Nasal prevention of SARS-CoV-2 infection by intranasal influenza-based boost vaccination

**DOI:** 10.1101/2021.10.21.465252

**Authors:** Runhong Zhou, Pui Wang, Yik-Chun Wong, Haoran Xu, Siu-Ying Lau, Li Liu, Bobo Wing-Yee Mok, Qiaoli Peng, Na Liu, Kin-Fai Woo, Shaofeng Deng, Rachel Chun-Yee Tam, Haode Huang, Anna Jinxia Zhang, Dongyan Zhou, Biao Zhou, Chun-Yin Chan, Zhenglong Du, Dawei Yang, Ka-Kit Au, Kwok-Yung Yuen, Honglin Chen, Zhiwei Chen

**Affiliations:** AIDS Institute, Li Ka Shing Faculty of Medicine, the University of Hong Kong; Pokfulam, Hong Kong Special Administrative Region, People’s Republic of China; Department of Microbiology, Li Ka Shing Faculty of Medicine, the University of Hong Kong; Pokfulam, Hong Kong Special Administrative Region, People’s Republic of China; State Key Laboratory for Emerging Infectious Diseases, the University of Hong Kong; Pokfulam, Hong Kong Special Administrative Region, People’s Republic of China; Centre for Virology, Vaccinology and Therapeutics Limited, the University of Hong Kong, Hong Kong Special Administrative Region, People’s Republic of China; Department of Clinical Microbiology and Infection Control, the University of Hong Kong-Shenzhen Hospital; Shenzhen, Guangdong, People’s Republic of China; National Clinical Research Center for Infectious Diseases, The Third People’s Hospital of Shenzhen and The Second Affiliated Hospital of Southern University of Science and Technology, Shenzhen, Guangdong, People’s Republic of China

**Author notes:** Correspondence: Zhiwei Chen (Lead contact), and Honglin Chen. Department of Microbiology, Li Ka Shing Faculty of Medicine, the University of Hong Kong, L5-45, 21 Sassoon Road, Pokfulam, Hong Kong SAR, People’s Republic of China. These authors made equal contributions (Runhong Zhou, Pui Wang, Yik-Chun Wong, Haoran Xu).

**Keywords:** SARS-CoV-2, Receptor binding domain, Mucosal immunity, Nasal prevention, PD1-based DNA vaccine, Live-attenuated influenza-based vaccine

## Abstract

**Background:** Vaccines in emergency use are efficacious against COVID-19, yet vaccine-induced prevention against nasal SARS-CoV-2 infection remains suboptimal.

**Methods:** Since mucosal immunity is critical for nasal prevention, we investigated an intramuscular PD1-based receptor-binding domain (RBD) DNA vaccine (PD1-RBD-DNA) and intranasal live attenuated influenza-based vaccines (LAIV-CA4-RBD and LAIV-HK68-RBD) against SARS-CoV-2.

**Findings:** Substantially higher systemic and mucosal immune responses, including bronchoalveolar lavage IgA/IgG and lung polyfunctional memory CD8 T cells, were induced by the heterologous PD1-RBD-DNA/LAIV-HK68-RBD as compared with other regimens. When vaccinated animals were challenged at the memory phase, prevention of robust SARS-CoV-2 infection in nasal turbinate was achieved primarily by the heterologous regimen besides consistent protection in lungs. The regimen-induced antibodies cross-neutralized variants of concerns. Furthermore, LAIV-CA4-RBD could boost the BioNTech vaccine for improved mucosal immunity.

**Interpretation:** Our results demonstrated that intranasal influenza-based boost vaccination is required for inducing mucosal and systemic immunity for effective SARS-CoV-2 prevention in both upper and lower respiratory systems.

**Funding:** This study was supported by the Research Grants Council Collaborative Research Fund (C7156-20G, C1134-20G and C5110-20G), General Research Fund (17107019) and Health and Medical Research Fund (19181052 and 19181012) in Hong Kong; Outbreak Response to Novel Coronavirus (COVID-19) by the Coalition for Epidemic Preparedness Innovations; Shenzhen Science and Technology Program (JSGG20200225151410198); the Health@InnoHK, Innovation and Technology Commission of Hong Kong; and National Program on Key Research Project of China (2020YFC0860600, 2020YFA0707500 and 2020YFA0707504); and donations from the Friends of Hope Education Fund. Z.C.’s team was also partly supported by the Theme-Based Research Scheme (T11-706/18-N).

## 1. Introduction

Since the discovery of coronavirus disease 2019 (COVID-19) in human clusters in December 2019 (1, 2), the causative agent, severe acute respiratory syndrome coronavirus 2 (SARS-CoV-2), has led to over 100 million infections globally with nearly 2.1 million deaths after just one year. While the COVID-19 pandemic continues to evolve, hundreds of vaccine candidates have been brought into preclinical and clinical trials at an expedited speed using various platforms of technology (3-5). Encouragingly, several vaccines have now acquired regulatory approvals for emergency use in various countries, yet few have been evaluated for inducing mucosal protection, especially for preventing robust SARS-CoV-2 infection in nasal turbinate (NT) (6, 7). NT in the upper respiratory tract (URT) is one of the most important portals of SARS-CoV-2 entry into humans. Ciliated nasal epithelial cells in NT have the highest expression of angiotensin-converting enzyme 2 (ACE2) and transmembrane serine protease 2 (TMPRSS2), supporting rapid and robust SARS-CoV-2 infection (8). Unlike SARS patients who have peak viral load in URT at day 10 after symptom onset (9), COVID-19 patients exhibit the highest viral loads in URT at or soon after the clinical presentation (1, 10, 11). Multiple SARS-CoV-2 variants can be transmitted simultaneously without a genetic bottleneck (12). Transmitted viruses then quickly impair host innate and adaptive immune responses (12), allowing for robust viral replication and asymptomatic viral spread. These findings indicated that the prevention of SARS-CoV-2 infection in NT is critical for pandemic control.

Current systemic vaccination and passive immunization are less effective or suboptimal for preventing SARS-CoV-2 in NT. Among vaccines authorized for emergency use, the Pfizer-BioNTech BNT162B2 mRNA, the Moderna mRNA-1273, the Oxford-ChAdOx1 and Novavax’s NVX-CoV2373 have released phase III results showing the efficacy of 95%, 94.1%, 70.4% and 89.3%, respectively, in preventing COVID-19 (13-15). The efficacy of the Oxford-ChAdOx1 vaccine against asymptomatic SARS-CoV-2 infection was only 3.8% among human vaccinees who received two standard doses (15). For passive immunotherapy, a receptor-binding domain (RBD)-specific human neutralizing antibody (NAb) LY-CoV555 has been evaluated in a phase 2 trial. Of three doses of 700 mg, 2800 mg and 7000 mg tested for monotherapy, only the medium dose of LY-CoV555 appeared to accelerate the natural decline of viral loads in nasopharyngeal swaps by day 11 (16). These results indicated that vaccine-induced or passive NAbs are suboptimal to prevent SARS-CoV-2 infection in human NT. Using the animal model, we reported that robust SARS-CoV-2 infection in NT may outcompete passive or vaccine-induced systemic NAbs, revealing a possible mechanism underlying the subprotection against asymptomatic infection (17). Considering that vaccine-induced subprotection may drive immune escape virus variants of concern and allow re-infection (18-20), we hypothesized that an effective vaccine should induce mucosal immunity for nasal protection to prevent SARS-CoV-2 asymptomatic infection and silent spread. Recently, thousands of vaccine-breakthrough infections in the United States have further indicated the need of priority research on nasal prevention of SARS-CoV-2 infection (21).

## 2. Methods

### 2.1 Mice

Male and female BALB/c and K18-hACE2 mice (aged 6–10 weeks) were obtained from the HKU Laboratory Animal Unit (LAU). The animals were kept in Biosafety Level-2 housing and given access to standard pellet feed and water *ad libitum* following LAU’s standard operational procedures (SOPs). The viral challenge experiments were then conducted in our Biosafety Level-3 animal facility following SOPs strictly.

### 2.2 Cell lines

HEK 293T-hACE2 and Vero E6 cells (RRID:CVCL_0574) were maintained in Dulbecco’s Modified Eagle Medium (DMEM) (Thermo Fisher Scientific) containing 10% fetal bovine serum, 2 mM L-glutamine and 100 U/mL penicillin and were incubated at 37□ in 5% CO2 setting (22).

### 2.3 Viruses

Confluent Vero-E6 cells were infected at 0.01 MOI with live SARS-CoV-2 HKU-13 strain (GenBank accession number MT835140). After 3 days incubation, virus supernatant was collected for titration by plaque assay using Vero-E6 cells.

### 2.4 Construction and Generation of PD1-based DNA and live attenuated influenza virus (LAIV)-based vaccine

Codon-optimized SARS-CoV-2 RBD gene was in fusion to a human soluble PD1 domain (PD1-RBD) using the pVAX plasmid as the backbone. To maintain functional domains of the fusion protein, a linker (GGGGS)3 was applied between the PD1 and RBD gene (23). The expression construct contained a human tissue plasminogen activator (tPA) secretory signal sequence to promote antigen secretion. The plasmid DNA transfection into HEK 293T cells was performed using polyethylenimine (PEI), and protein expression was detected by Western blot. The pHW2000-DelNS1-RBD plasmid was constructed by inserting the tPA-linked RBD between the noncoding region (NCR) and autoproteolytic cleavage site (2A) in the pHW2000-DelNS1 plasmid. The V5 tag was added to the C terminal of RBD for better detection of RBD. To rescue the virus, eight pHW2000 plasmids containing the DelNS1-RBD and the other 7 influenza virus genomic segments, together with an NS1 expression plasmid, were transfected into 293T cells using Transit-LT1 (Mirus) according to the manufacturer protocol. After overnight incubation at 33 □, DNA mix was removed, and MEM supplemented with 1 μg/ml N-tosyl-L-phenylalanine chloromethyl ketone (TPCK)-treated trypsin (Sigma) was added. Virus supernatant was collected 72 hours later and designated passage 0 (p0) virus and was further passaged in chicken embryonated eggs for 48 hours at 33 □. Viruses were aliquoted and titrated by plaque assay using MDCK cells. Two intranasal recombinant LAIV DelNS1-RBD vaccine strains, namely LAIV-CA4-RBD and LAIV-HK68-RBD, were generated using A/CA/04/2009 (H1N1) and A/Hong Kong/1/68 (H3N2) surface proteins (HA and NA) in the A/CA/04/2009 (H1N1) DelNS1 backbone by a reverse genetic procedure (24).

### 2.5 Animal immunization and SARS-CoV-2 challenges

All animal experiments were approved by the Committee on the Use of Live Animals in Teaching and Research of the University of Hong Kong (HKU). For vaccine immunization, 6-8-weeks old BALB/c or K18-ACE2 transgenic mice received DNA immunization by intramuscular/electroporation (i.m./EP) with 50 μg PD1-RBD-DNA. The voltage of EP was pre-set 60 V in the TERESA DNA Delivery Device (Shanghai Teresa Healthcare Sci-Tech Co., Ltd). Mice intramuscularly received 1/5 clinical doses of Pfizer/BioNTech or Sinovac vaccine. Mice received LAIV-CA4-RBD or LAIV-HK68-RBD immunization by intranasal inoculation of 10^6^ PFU per mouse at a 3-week interval under anesthesia. Blood sera was collected for anti-RBD IgG and neutralization detection. Mice were sacrificed and cells from lungs and spleen were harvested and subjected to intracellular cytokine staining (ICS) assay. For the SARS-CoV-2 challenge, BALB/c mice were anesthetized and transduced intranasally with 4×10^8^ FFU of Ad5-hACE2, kindly provided by Dr. Jincun Zhao (25), in 70 μl DMEM. The transduced BALB/c mice or K18-ACE2 transgenic mice were intranasally infected with live wild type SARS-CoV-2 (HKU clone 13) at a dose of 1×10^4^ PFU. Infected animals were sacrificed for endpoint analysis at day 4 post infection (4 dpi). All animal experiments related to SARS-CoV-2 were performed in a biosafety level 3 laboratory in HKU.

### 2.6 Enzyme-linked immunosorbent assay (ELISA)

ELISA was performed to detect SARS-CoV-2 RBD-specific IgG, as previously described (26). In brief, 96-well plates (Costar) were coated with recombinant SARS-CoV-2 RBD antigen (25 ng/well, Sino Biological) at 4°C overnight. After washing with PBST (0.05% Tween-20 in PBS), the plates were blocked with 4% skim milk in PBS for 1 hour at 37°C and incubated with serially diluted patient plasma for 1 hour at 37°C. After washing with PBST, goat anti-mouse IgG or anti-mouse IgA conjugated with HRP (Invitrogen) was added, and the whole solution was incubated for 1 hour, followed by washing and the addition of 50 μl HRP chromogenic substrate 3,3’,5,5’-TMB (Sigma). Optical density (OD) values were measured at 450 nm using the VARIOSKANTM LUX multimode microplate reader (Thermo Fisher Scientific). Area under the curve (AUC) was measured using GraphPad Prism v8, setting the baseline with the defined endpoint (average of negative control wells + 10 standard deviation) and taking the total peak area as previous described (27).

### 2.7 Pseudotyped viral neutralization assay

To determine the neutralizing activity of mouse sera and BAL, specimens were inactivated at 56□ for 30 min before the pseudotype viral entry assay as previously described (22, 28). The results of this assay correlated strongly with that of neutralization assay using replication-competent SARS-CoV or SARS-CoV-2 (8, 29). The plasmids encoding for D614G, Alpha, Beta and Delta variants were kindly provided by Dr. David D. Ho. In brief, different SARS-CoV-2 pseudotype viruses were generated through co-transfection of 293T cells with 2 plasmids, pSARS-CoV-2 S and pNL4-3Luc_Env_Vpr, carrying the optimized SARS-CoV-2 S gene and a human immunodeficiency virus type 1 backbone, respectively. At 48 hours post-transfection, viral supernatant was collected and frozen at -80°C. Serially diluted serum samples were incubated with 200 TCID50 of pseudovirus at 37°C for 1 hour. The serum-virus mixtures were then added into pre-seeded HEK 293T-hACE2 cells. After 48 hours, infected cells were lysed, and luciferase activity was measured using Luciferase Assay System kits (Promega) in a Victor3-1420 Multilabel Counter (PerkinElmer). The 50% inhibitory concentrations (IC50) of each specimen were calculated using non-linear regression in GraphPad Prism v8 to reflect anti-SARS-CoV-2 potency.

### 2.8 Surface and intracellular cytokine staining (ICS)

The lung cells of mice were washed one time with staining buffer (PBS contained 2% FBS) followed by staining with anti-mouse antibodies for 30 min at 4 □, including dead cell dye (Zombie Aqua, Biolegend Cat# 423102), CD19-FITC (Biolegend Cat# 152404, RRID: AB_2629813), CD11b-PerCP/Cy5.5 (Biolegend Cat# 101228, RRID: AB_893232), CD11c-PE-Cy7 (Biolegend Cat# 117318, RRID: AB_493568), Ly6c-APC-Fire750 (BioLegend Cat# 128046, RRID: AB_2616731), F4/80-BV421 (Biolegend Cat# 123137, RRID: AB_2563102), Ly6G-PE (BioLegend Cat# 127608, RRID: AB_1186099), CD103-BV785 (Biolegend Cat# 121439, RRID: AB_2800588) and I-A/I-E-BV605 (Biolegend Cat# 107639, RRID: AB_2565894). To measure antigen-specific T cell response, lymphocytes from mouse lung and spleen were stimulated with 1 μg/mL SARS-CoV-2 RBD peptide pool (15-mer overlapping by 11, spanning the whole RBD sequence at Spike_306-543_). Cells were incubated at 37□ overnight, and BFA was added at 2 hours post-incubation. PMA/ionomycin stimulation was included as the positive control. After overnight incubation, cells were washed with staining buffer (PBS containing 2% FBS) and surface stained with anti-mouse-CD4-PerCP/Cy5.5 (Biolegend Cat# 116012, RRID: AB_2563023), anti-mouse-CD8-BV785 (Biolegend Cat# 100750, RRID: AB_2562610), anti-mouse CD69-BV711 (Biolegend Cat# 104537, RRID: AB_2566120) and anti-mouse CD103-BV421 (Biolegend Cat# 121422, RRID: AB_2562901). Zombie aqua staining was used to exclude dead cells. For intracellular staining, cells were fixed and permeabilized with BD Cytofix/Cytoperm (BD Biosciences) prior to staining with anti-mouse-IFN-γ-APC (Biolegend Cat# 505810, RRID: AB_315404), anti-mouse-TNF-α-PE (Biolegend Cat# 506306, RRID: AB_315427) and anti-mouse-IL-2-PE-Cy7 (Biolegend Cat# 503832, RRID: AB_2561750). Stained cells were acquired by FACSAriaIII Flow Cytometer (BD Biosciences) and analyzed with FlowJo software (v10.6) (BD Bioscience).

### 2.9 Viral RNA quantification

Half nasal turbinates and lung tissues were homogenized and subjected to viral load determination by quantitative SARS-CoV-2-specific RdRp/Hel reverse-transcription polymerase chain reaction assay (10). Total RNA was extracted using RNAeasy mini kit (Qiagen) and followed by reverse transcription (PrimeScript II 1^st^ Strand cDNA Synthesis Kit). The real-time RT-PCR assay for SARS-CoV-2-RdRp/Hel RNA and NP subgenomic RNA detection was performed using One-Step TB Green PrimeScript RT-PCR Kit II according to the manufacture’s instruction. Mouse β-actin was used as normalization.

### 2.10 Plaque assay

Infectious virus titration was determined by plaque assay. Confluent Vero-E6 cells in a 12-well plate were incubated with 10-fold serially diluted tissue homogenates for 1 h. The virus supernatant was discarded, and cells were then overlaid with 1% agarose in DMEM and further incubated for 3 days at 37 □ followed by overnight fixation via 4% PFA. Agarose gels were removed, and plaques were visualized by 1 % crystal violet.

### 2.11 Histopathology and Immunofluorescence (IF) Staining

Tissues collected at necropsy were fixed in zinc formalin and then processed into paraffin-embedded tissue blocks. The tissue sections (4 μm) were stained with hematoxylin and eosin (H&E) for light microscopy examination. For identification and localization of SARS-CoV-2 nucleocapsid protein (NP) in organ tissues, IF staining was performed on deparaffinized and rehydrated tissue sections using rabbit anti-SARS-CoV-2-N protein antibody together with AF568-conjugated goat anti-rabbit IgG (Invitrogen). Briefly, the tissue sections were first treated with antigen unmasking solution (Vector Laboratories) in a pressure cooker. After blocking with 0.1% Sudan black B for 15 min and 1% bovine serum albumin (BSA)/PBS at RT for 30 min, the primary antibody rabbit anti-SARS-CoV-2-N antibody (1:4000 dilution with 1% BSA/PBS) was incubated at 4°C overnight. This step was followed by AF568-conjugated goat anti-rabbit IgG for 30 min and then mounted with 4’,6-diamidino-2-phenylindole (DAPI). All tissue sections were examined, the images were captured with a Carl Zeiss LSM780 confocal microscope, and the mean fluorescence intensity (MFI) was further measured by ImageJ v1.53c.

### 2.12 Statistical analysis

Statistical analysis was performed with the GraphPad Prism 8 Software. Data represent mean values or mean values with SEM. Significant differences between the means of multiple groups were tested using a one-way analysis of variance (ANOVA) followed by Tukey’s multiple comparisons test. Significant differences between the two groups were performed using the 2-tailed Student’s t-test. P < 0.05 was considered statistically significant.

### 2.13 Animal study approval

All experimental procedures were approved by the Committee on the Use of Live Animals in Teaching and Research (CULATR 5350-20) of the University of Hong Kong.

## 3. Results

### 3.1 Construction and characterization of PD1-based DNA and influenza-based vaccines

To address the hypothesis, we sought to study an intramuscular program death 1 (PD1)-based RBD DNA vaccine (PD1-RBD-DNA) and two intranasal live attenuated influenza virus (LAIV)-based vaccines (LAIV-HK68-RBD and LAIV-CA4-RBD). Recently, Y Liu et al. demonstrated that antibodies against the N-terminal domain (NTD) could enhance the binding capacity of the spike protein to ACE2 and infectivity of SARS-CoV-2 (30). To avoid the potential full spike (S)-associated side effects (22, 30, 31), we chose RBD as the vaccine immunogen. Since delayed cytotoxic T lymphocyte (CTL) responses are likely associated with COVID-19 severity (12), we constructed the PD1-RBD-DNA vaccine (Fig. S1A) that might elicit enhanced antibody and CD8 T cell responses (23). The expression of soluble PD1-RBD protein (∼80 kDa) was readily detected in supernatants of transfected HEK293T cells by Western blot analysis using either anti-SARS-CoV-2 RBD antibody (green lane) or anti-human PD-1 (red lane) antibody (Fig. S1B). The released PD1-RBD was able to bind PD-L (Fig. S1C), which might help antigen targeting to dendritic cells for cross-presentation (23). Meantime, the NS1-deleted (DelNS1) LAIV was engineered to express the same RBD (Fig. S1D), aiming to induce mucosal NAb and T cell immune responses (32). We previously characterized a panel of DelNS1 influenza viruses and evaluated their potential for being used as vaccine vectors (32). Two intranasal recombinant LAIV DelNS1-RBD vaccine strains, namely LAIV-CA4-RBD and LAIV-HK68-RBD, were generated using A/CA/04/2009 (H1N1) and A/Hong Kong/1/68 (H3N2) surface proteins (HA and NA) in the A/CA/04/2009 (H1N1) DelNS1 backbone by a reverse genetic procedure (24). The SARS-CoV-2 RBD and LAIV NP proteins were stably expressed in MDCK cells by Western blot after 5 times of viral passages in MDCK cells (Fig. S1E).

### 3.2 Systemic and mucosal antibody responses of vaccine regimens

We then chose the immune competent SARS-CoV-2/BALB/c mouse model (25), which was based on available antibody reagents to understand the potential correlate of immune protection. As compared with the vector control group (v1) that received 50 μg intramuscular electroporation (i.m./EP) pVAX plasmid prime plus i.n. 10^6^ PFU LAIV-68 vector boost (Fig. 1A), we tested groups treated by the homologous 50 μg i.m./EP PD1-RBD-DNA twice (v2), a heterologous i.n. 10^6^ PFU LAIV-CA4-RBD prime plus i.n. 10^6^ PFU LAIV-HK68-RBD boost (v3), a heterologous 50 μg i.m./EP PD1-RBD-DNA prime plus i.n. 10^6^ PFU LAIV-68-RBD boost (v4) and a heterologous i.n. 10^6^ PFU LAIV-CA4-RBD prime plus 50 μg i.m./EP PD1-RBD-DNA boost (v5) at 3-week intervals, consistent with COVID-19 vaccines under emergency use. Vaccine-induced antibody responses were determined at day 9 (acute phase) and day 69 (memory phase) post the second vaccination. Both peripheral blood and bronchoalveolar lavage (BAL) samples were collected from vaccinated mice for antibody detection by ELISA and pseudovirus neutralizing assays. We found that the PD1-RBD-DNA/LAIV-HK68-RBD regimen (v4) elicited and significantly sustained the highest amounts of RBD-specific IgG (acute: mean 5.2, range 4.86-5.45 logs AUC; memory: mean 4.62, range 4.54-4.69 logs AUC) and NAbs (acute: mean 4.19, 4.04-4.34 logs IC_50_; memory: mean 2.89, 2.4-3.23 logs IC_50_) in sera during both acute and memory phases as compared with other groups (Fig. 1B-C). The RBD-specific IgG titer and NAb IC_50_ values were correlated positively (Fig. 1D), similar to COVID-19 patients’ sera (8). Moreover, v4 animals also developed and sustained significantly higher amounts of RBD-specific mucosal IgG and IgA in BAL during both acute and memory phases as compared with other groups (Fig. 1E-F). The amount of BAL NAbs in v4 mice were at mean 2.80 (range 2.22-2.87) and mean 2.59 (range 2.02-3.15) logs IC_50_ at the acute and memory phase, respectively (Fig. 1G). In contrast, v2 and v5 elicited similar amounts of NAb to v4 at the acute phase, yet these responses did not sustain into the memory phase. Moreover, despite heterologous intranasal immunizations twice, the v3 regimen did not induce equally potent and sustained mucosal IgG and IgA as well as NAb responses, as compared with the v4 group (Fig. 1G). Both BAL IgG and IgA titers correlated positively with the BAL NAb values (Fig. 1H) despite of the higher amount and better acute/memory of BAL IgG than BAL IgA. Interestingly, serum NAb IC_50_ values were correlated positively with the BAL NAbs IC_50_ values at the memory phase, but not correlated at the acute phase (Fig. 1I). These results indicated that the v4 regimen is likely unique for inducing potent and sustained systemic and mucosal memory IgG/IgA NAb responses.

**Fig. 1.**
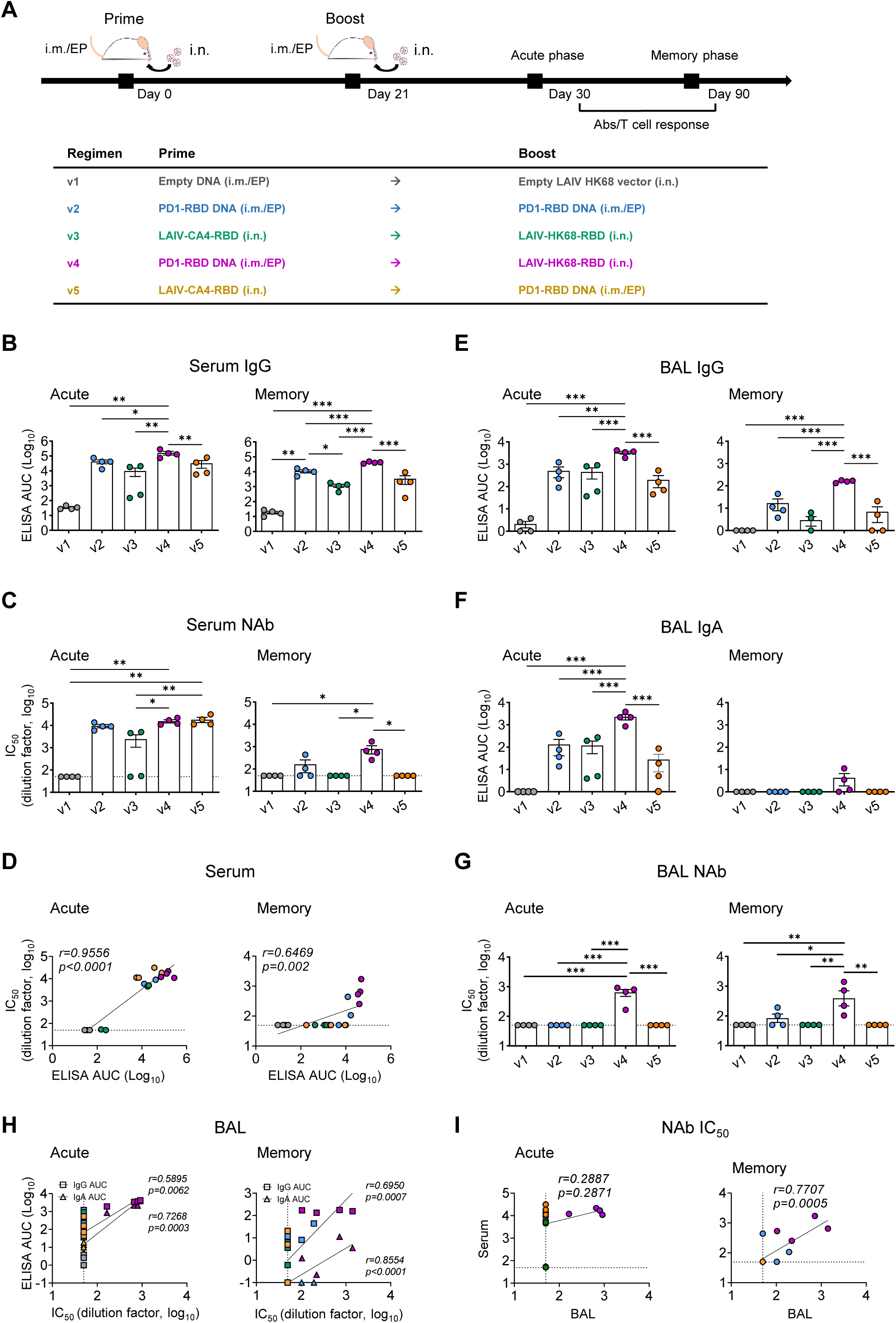
Vaccine-induced systemic and mucosal antibody responses. (**A**) Vaccine immunization schedule and grouping for BALB/c mice (n=8/group). Blood and bronchoalveolar lavage (BAL) were collected and subjected to antibody response analysis at 9 (acute phase, n=4/group) or 69 (memory phase, n=4/group) days post the 2nd immunization, respectively. In sera, (**B**) RBD-specific IgG titer, (**C**) NAb IC_50_ values and (**D**) positive correlations between RBD-specific IgG titer and NAb IC50 values were analyzed for both acute and memory phases. In BAL, (**E**) RBD-specific IgG titer, (**F**) RBD-specific IgA titer, (**G**) NAb IC_50_ values and (**H**) positive correlations between RBD-specific IgG (square) or IgA titers (triangle) and BAL NAb IC_50_ values were analyzed for both acute and memory phases. (**I**) Correlate analysis between Peripheral NAb IC_50_ values and BAL NAb IC50 values at both acute and memory phase. RBD-specific IgG or IgA titers were determined by ELISA at serial dilutions. The area under the curve (AUC) represented the total peak area calculated from ELISA OD values. Neutralization IC_50_values were determined against SARS-CoV-2-Spike-pseudovirus infection of 293T-huACE2 cells. Correlation analysis was performed by linear regression using GraphPad Prism 8.0. Each symbol represents an individual mouse with color-coding for corresponding groups. Error bars indicate the standard error of the mean. Statistics were generated using one-way ANOVA followed by Tukey’s multiple comparisons test. *p<0.05; **p<0.01.

### 3.3 Acute and memory T cell responses of vaccine regimens

Since SARS-CoV-2-specific T cell responses are essential for control and resolution of viral infection (12, 33), we sacrificed five groups of animals to measure vaccine-induced T cell immune responses at day 9 (acute phase) and day 69 (memory phase) post the second vaccination. Lymphocytes were isolated from both lungs (effector site) and spleens (secondary lymph organ) of vaccinated mice for comparison. We found that the v4 regimen elicited and sustained significantly higher frequencies of RBD-specific IFN-γ^+^ CD8 T cells in lungs and spleens during both acute (Fig. 2A-B, Fig. S2A-B) and memory (Fig. 2D-E, Fig. S2D-E) phases as compared with other groups. Similar trends were found with IFN-γ^+^ CD4 T cells elicited in the v4 regimen but at lower frequencies. At the acute phase, the v4 regimen elicited the highest mean frequency of RBD-specific IFN-γ^+^ CD8 T cells (mean 28.83%, range 22-34.8%) in lungs (Fig. 2B), which was even higher than that in spleens (mean 5.54%, range 3.63-7.06%) (Fig. S2B). These cells included the highest frequencies of polyfunctional CTLs with a capacity of releasing two (mean 21.09%, range 13.87-25.31%) or three (mean 2.27%, range 1.02-3%) cytokines (Fig. 2C), which was also higher than those in splenic CD8 T cells releasing two (mean 6.68%, range 4.68-9.17%) or three cytokines (mean 0.92%, range 0.7-1.36%) (Fig. S2C). At the memory phase, the v4 regimen sustained the highest mean frequency of RBD-specific IFN-γ^+^ CD8 T cells in lungs (mean 6.11%, range 2.05-9.7%) (Fig. 2E) and spleens (mean 2.35%, range 0.7-4.78%) (Fig. S2E) as compared with other groups. These cells included the highest frequencies of polyfunctional CTLs with a capacity of releasing two (mean 8.66%, range 2.61-12.41%) or three (mean 2.2, range 0.66-3.09%) cytokines (Fig. 2F), which was higher than those in splenic CD8 T cells releasing two (mean 3.59%, range 1.28-6.83%) or three cytokines (mean 0.69%, range 0.31-1.17%) (Fig. S2F). These results demonstrated that besides Nabs, the v4 regimen also induced potent and polyfunctional memory CD8 T cell responses, especially in lungs. Since overall immune responses induced by the heterologous v3 regimen were much weaker than those by the v4 regimen, we also exanimated T cell responses against influenza immunodominant nucleoprotein (NP) (34). At acute phase (Fig. S3A), the v3 regimen induced the highest frequencies of CD8 T cell response against influenza NP in lungs (mean 19.55%, range 16.2-21.7%) as compared with v1 (mean 8.3%, range 7.77-8.9%) and v5 (mean 3.99%, range 2.55-5.91%). Similar results were observed at the memory phase (Fig. S3B). In contrast, the v4 regimen induced significantly lower influenza NP-specific T cell response at both acute (mean 2.26%, range 1.68-2.58%) and memory (mean 0.77%, range 0.49-1.06%) phases. The heterologous prime using PD1-RBD-DNA instead of LAIV-CA4-RBD, therefore, offered an advantage in promoting the RBD immunodominance likely by avoiding anti-vector immune responses.

**Fig. 2.**
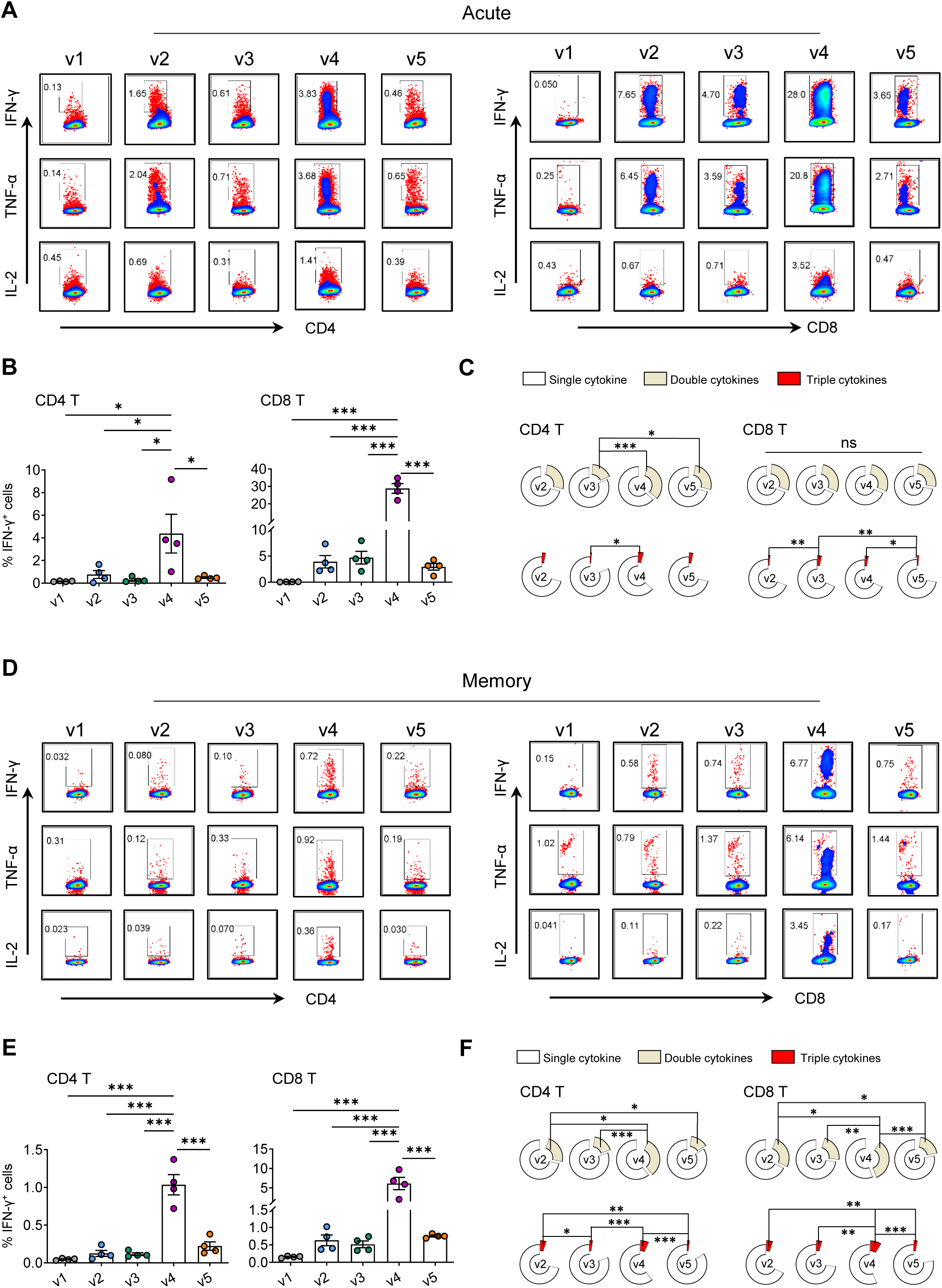
Vaccine-induced acute and memory T cell responses in lungs. The vaccine immunization schedule for BALB/c mice was the same as described in Fig. 1A. Cells from lungs were collected and subjected to SARS-CoV-2 RBD-specific T cell responses by *ex vivo* RBD peptide pool stimulation followed by ICS at both acute (**A-C**) and memory (**D-F**) phase. (**A, D**) Representative dot plots present the gating of IFN-γ^+^, TNF-α^+^ or IL-2^+^ CD4 T (left) and CD8 T (right) against SARS-CoV-2 RBD. (**B, E**) Quantified results depict the percentage of IFN-γ^+^ CD4 T (left) and IFN-γ^+^ CD8 T (right). Each symbol represents an individual mouse. Error bars indicate the standard error of the mean. (**C, F**) The pie charts indicate the proportion of single or double or triple cytokines produced by CD4 T (left) and CD8 T (right). Statistics were generated using one-way ANOVA followed by Tukey’s multiple comparisons test. *p<0.05; **p<0.01; ***p<0.001.

### 3.4 Protective efficacy against intranasal SARS-CoV-2 infection

To investigate the efficacy of various vaccine regimens against the live intranasal SARS-CoV-2 challenge, we subsequently immunized additional groups of BALB/c mice (n=6 per group) using the same doses and time interval as described above (Fig. 3A). We did not include v5 due to low mucosal immunogenicity and limited space in our animal P3 facility. Sera were collected at day 9 and day 28 after the 2^nd^ immunization to monitor anti-RBD IgG (Fig. 3B) and neutralization (Fig. 3C). Consistently, the highest IgG (acute: mean 4.79, range 4.7-4.89 logs AUC; memory: mean 2.6, range 2.03-2.81 logs AUC) and neutralizing (acute: mean 3.9, range 3.69-4.14 logs IC_50_; memory: mean 2.94, range 2.34-3.33 logs IC_50_) titers were induced in mice by the v4 regimen at both acute and memory phases. Immunized mice were then transduced with Ad5-hACE2 at the memory phase, 29 days post the boost vaccination for expressing human ACE2 in nasal turbinate and lung (Fig. 3D), followed by the intranasal SARS-CoV-2 challenge 6 days later as previously described (25). At day 4 after the viral challenge, mice were sacrificed for analysis. Lung specimen was harvested to quantify infectious viruses by plaque assay, viral load by real-time PCR (RT-PCR) and infected cells by immunofluorescence staining (IF). We found that all vaccinated animals had decreased infectious plaque-forming units (PFU) to the limit of detection (10 PFU/mL) in lungs (Fig. 3E). The v4 regimen, however, resulted in the most significant genomic RdRp (gRdRp) drop in lungs by an average of 2.31 logs compared with 1.81 logs in v2 mice and 1.62 logs in v3 mice (Fig. 3F). A similar observation was found in the measurement of nucleocapsid protein (NP) subgenomic RNA (sgNP) (Fig. 3G). These findings demonstrated that immune responses induced by v2, v3 and v4 regimens had achieved significant protection in lungs. To determine viral infection in both upper and lower respiratory systems, we further performed immunofluorescence staining of SARS-CoV-2 NP antigen in both lung (Fig. 3H) and NT (Fig. 3I) tissues. Since murine NT was too small to be sliced for viral load tests, we only used it for the NP staining to maintain the necessary tissue structure. While significantly reduced NP^+^ cells were observed in lungs of v2 and v3 mice, infected cells were barely found in lungs of v4 mice. Furthermore, no significantly reduced NP^+^ cells were found in NT of v2 and v3 mice as compared with v1 mice, but only a few NP^+^ cells were seen in v4 mice. Our results demonstrated that while protection was consistently found in lungs of vaccinated animals, significant prevention of robust SARS-CoV-2 infection in NT was only achieved by the v4 regimen.

**Fig. 3.**
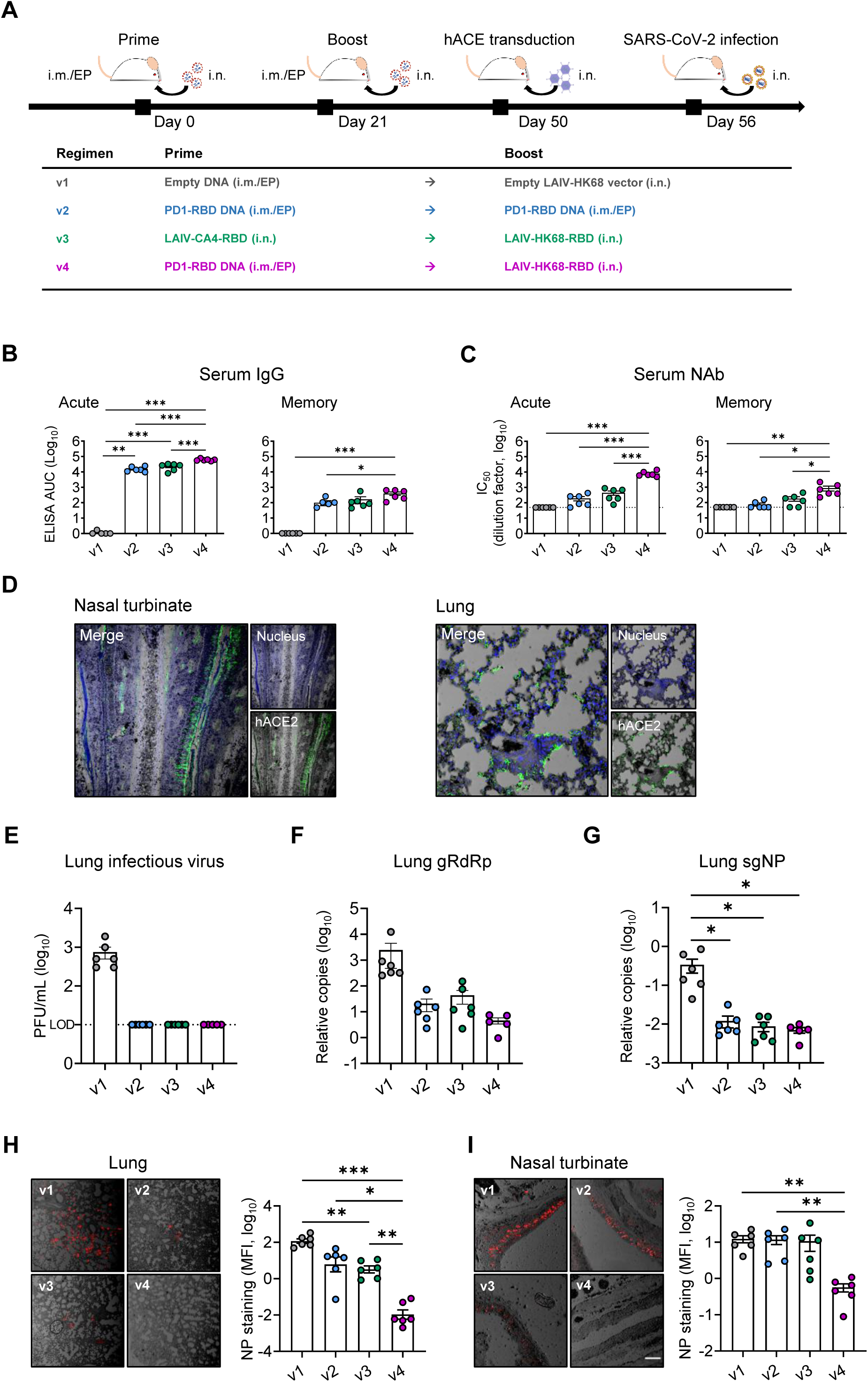
Protective efficacy of vaccine regimens in huACE2-transduced mice. (**A**) Experimental schedule and grouping of BALB/c mice (6 mice/group). At day 29 post the 2^nd^ immunization, mice were transduced to express human ACE2 in vivo by inoculating 4×10^8^ plaque-forming units (PFU) Ad5-hACE2 intranasally. Six days later, mice were challenged intranasally with 10^5^ PFU SARS-CoV-2 and sacrificed at day 4 post-infection. (**B, C**) Serum samples were collected for detection of anti-RBD IgG (**B**) and neutralizing antibody (**C**) against pseudovirus, respectively, at day 9 (acute, left) and day 28 (memory, right) post the 2^nd^ immunization. The Area under the curve (AUC) represents the total peak area calculated from ELISA OD values. (**D**) Human ACE2 expression (green) after the Ad5-hACE2 transduction was evaluated in NT and lung by immunofluorescence (IF) staining using the hACE2-specific antibody at day-6 post-transduction. With nuclei counter staining (blue), images were shown at the 10× magnification. (**E**) A viral plaque assay was used to quantify infectious viruses in lung homogenates. Log_10_-transformed plaque-forming units (PFU) per mL of tissue extractions were shown for each group. LOD: limit of detection. (**F, G**) Sensitive RT PCR was used to quantify SARS-CoV-2 RdRp RNA (**F**) and NP subgenomic RNA (**G**) copy numbers (normalized by β-actin) in lung homogenates. Confocal images showed SARS-CoV-2 NP positive (red) cells (20×) in lungs (**H**) and NT (**I**) in the bright field. Mean fluorescent intensities (MFI) of NP^+^ cells in lung and NT were measured using ImageJ software and plotted with GraphPad prism. Each symbol represents an individual mouse with consistent color-coding. Error bars indicate the standard error of the mean. Statistics were generated using one-way ANOVA followed by Tukey’s multiple comparisons test. *p<0.05; **p<0.01; ***p<0.001.

### 3.5 Infection-recalled NAb for correlate of protection

Although we were not allowed to bring tissue specimens for measuring T cell immunity outside the animal P3 laboratory after the SARS-CoV-2 challenge, we managed to determine if viral infection recalled vaccine-induced NAbs for viral neutralization and clearance (35). By testing RBD-specific IgG and NAb at day 28 (before challenge) and day 39 (also 4 dpi) after the 2^nd^ vaccination, we found that SARS-CoV-2 infection indeed recalled significantly anti-RBD IgG responses in all v2, v3 and v4 animals (Fig. S4A). Notably, 62.2-fold and 65.7-fold higher amounts of recalled NAb were found by the v4 regimen (v4, mean 4.64, range 3.22-5.03 logs IC_50_) than the v2 and v3, respectively (Fig. S4B). Furthermore, there were significant negative correlations between viral loads and IgG or NAb IC_50_ values (Fig. S4C-D), as well as between NP^+^ cells and IgG or NAb IC_50_ values in lungs (Fig. S4E-F) and in NT (Fig. S4G-H). Moreover, since the T_H_1 immune response is likely associated with protective immune responses against COVID-19 (36), we also evaluated the ratios of IgG1 and IgG2a in all vaccinated mice. The v4 regimen likely induced a T_H_1 bias with a higher IgG2a/IgG1 ratio (mean 59.57, range 1.44-279.2) than those induced by v2 (mean 6.72, range 2.63-15.14) and v3 (mean 7.65, range 4.25-12.72) (Fig. S4I). Our results demonstrated that v4 vaccine-induced high amounts of NAbs correlated with protective efficacy in both NT and lung.

### 3.6 SARS-CoV-2 prevention in both upper and lower respiratory tracts of K18-hACE2 mice

To further determine the vaccine efficacy, we tested the PD1-RBD-DNA/LAIV-HK68-RBD regimen in K18-hACE2 transgenic mice, one of commonly used animal models for studying COVID-19 (37-39). 8-week-old K18-hACE2 mice were vaccinated with various regimens using the doses and time interval as described above (Fig. 4A). The titer of RBD-specific IgG (Fig. 4B) and neutralization antibody (Fig. 4C) was measured in serum at day-9 and day-33 (acute phase) after the 2^nd^ immunization. Consistently, the PD1-RBD-DNA/LAIV-HK68-RBD regimen (v4) induced and sustained the highest amount of RBD-specific serum IgG (acute: mean 4.38, range 3.81-4.84 logs AUC; memory: mean 4.79, range 4.27-5.25 logs AUC) and Nabs (acute: mean 3.74, range 3.10-4.30 logs IC_50_; memory: mean 4.02, range 2.48-4.64 logs IC_50_) during both acute and memory phases as compared with other groups. At day 38 after the 2^nd^ immunization, mice were intranasally challenged with 10^4^ PFU of SARS-CoV-2 (HKU Clone 13). In lungs, no measurable infectious viruses (detection limit: 10 PFU/mL) were found in all v4 mice and 40% v2 mice, whereas infectious viruses were detected in all v1 and v3 mice (Fig. 4D). The more sensitive RT-PCR further demonstrated that the v4 regimen resulted in the significant gRdRp drop by an average of 3.57 logs as compared with 0.81 logs in v2 mice and 0.25 logs in v3 mice (Fig. 4E) at 4 dpi. A similar observation was found with the measurement of NP subgenomic RNA (Fig. 4F). Importantly, no measurable infectious viruses were detected in nasal turbinate of v4 mice (Fig. 4G) and the v4 regimen significantly suppressed viral genomic/subgenomic RNA (gRdRp and sgNP) (Fig. 4H-I). Further IF staining of NP antigen confirmed that significantly reduced NP^+^ cells were observed in both lungs (Fig. 4K) and nasal turbinates (Fig. 4L) of v4 mice as compared with other regimens. Daily monitoring of body weight after challenge showed that SARS-CoV-2 infection caused average 15%, 8% and 6% weight loss in v1, v2 and v3 mice at 4 dpi, respectively. In contrast, no weight loss was found in v4 mice during the experimental period (Fig. 4J). To further characterize the injury of lung and nasal turbinate, we performed pathological analysis on specimens by haematoxylin and eosin (H&E)-staining. Acute lung injury with peribronchiolar infiltration, alveolar space infiltration and exudation were visualized in v1 mice, while v2 and v3 mice showed less alveolar space infiltration. In contrast, only mild peribronchiolar infiltration was observed in v4 mice (Fig. 4M). In nasal turbinate, damage of epithelium with extensive submucosal immune cell infiltration was observed in v1, v2 and v3 mice, but not in v4 mice (Fig. 4N). Consistent results were found in the flow cytometry analysis of immune cell compositions in lungs. A significant decrease in frequency of alveolar macrophages as well as reduced CD103^+^ DCs and CD11b^+^ DCs (Fig. S5A-B) was found in the v4 mice as compared with other groups. These results in K18-hACE2 mice, therefore, further demonstrated that the PD1-RBD-DNA/LAIV-HK68-RBD regimen prevents robust SARS-CoV-2 infection not only in lung and but also in NT with minimal infection-associated inflammation and injury.

**Fig. 4.**
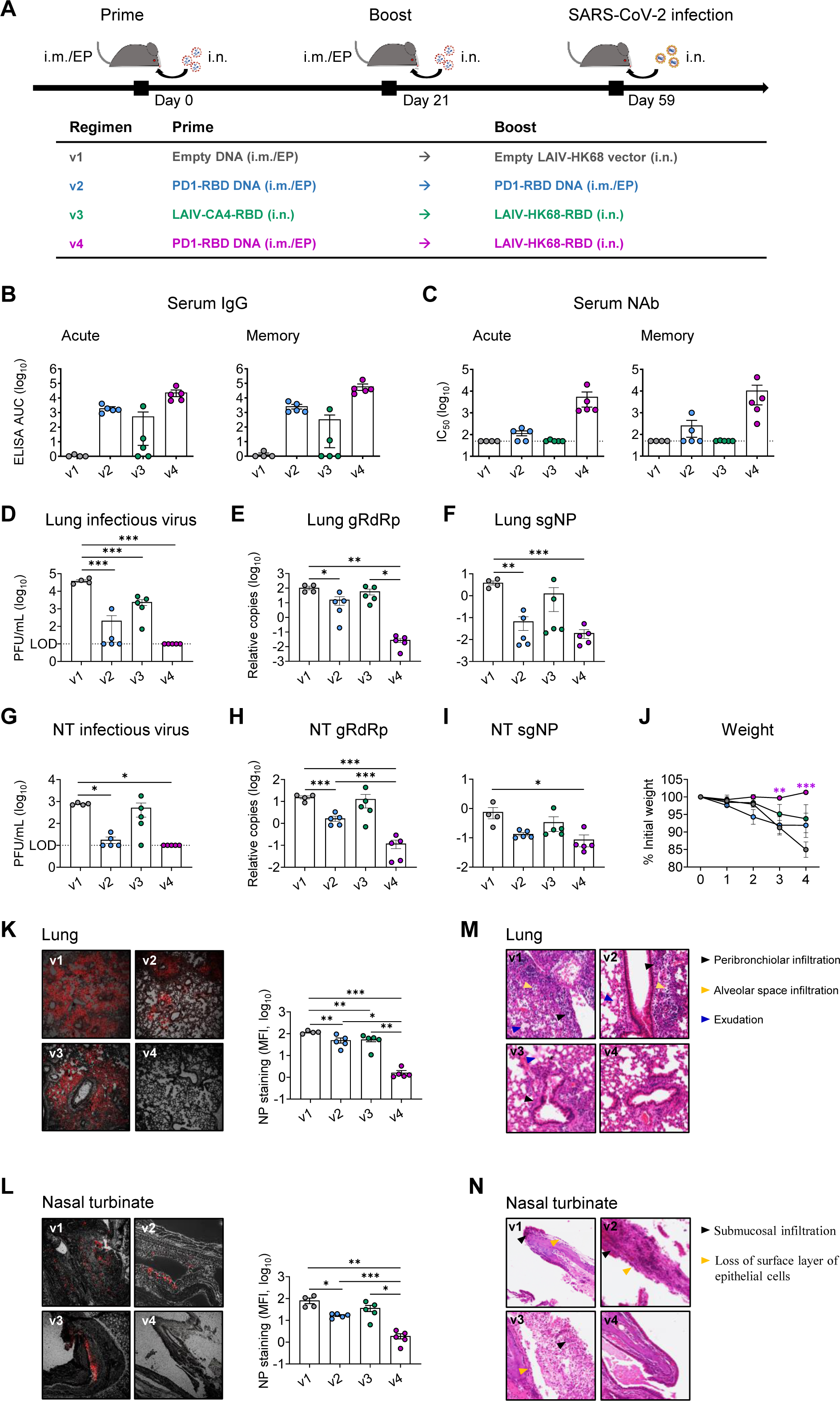
PD1-RBD-DNA/LAIV-HK68-RBD regime prevents SARS-CoV-2 infection in the upper and lower respiratory tracts of hACE2 transgenic mice. **(A)** Experimental schedule and grouping of K18-hACE-2 mice (4 mice in v1 group and 5 mice in other groups). (**B, C**) Serum samples were collected for detection of anti-RBD IgG (**B**) and neutralizing antibody (**C**) against pseudovirus, respectively, at day 9 (acute, left) and day 33 (memory, right) post the 2^nd^ immunization. The Area under the curve (AUC) represents the total peak area calculated from ELISA OD values. A viral plaque assay was used to quantify infectious viruses in lung (**D**) and NT (**G**) homogenates. Log_10_-transformed plaque-forming units (PFU) per mL of tissue extractions were shown for each group. LOD: limit of detection. Sensitive RT PCR was used to quantify SARS-CoV-2 RdRp RNA (**E, H**) and NP subgenomic RNA (**F, I**) copy numbers (normalized by β-actin) in lung and NT homogenates. (**J**) Daily body weight was measured after infection. Differences between groups that were given the different vaccine regime versus PBS were determined using a 2-tailed Student’s t test. Confocal images showed SARS-CoV-2 NP positive (red) cells in lungs (10×) (**K**) and NT (20×) (**L**) in the bright field. Mean fluorescent intensities (MFI) of NP^+^ cells in lung and NT were measured using ImageJ software and plotted with GraphPad prism. Representative images of animal lung (**M**) and NT (**N**) tissues by H&E (20×). Each symbol represents an individual mouse with consistent color-coding. Error bars indicate the standard error of the mean. Statistics were generated using one-way ANOVA followed by Tukey’s multiple comparisons test. *p<0.05; **p<0.01; ***p<0.001.

### 3.7 Both NAb and antigen-specific CD8 T cells associated with virus control in K18-hACE2 mice

Compare to the serum collected at day 5 before the challenge, the titer of RBD-specific IgG (Fig. 5A) and NAb (Fig. 5B) did not increase in sera collected at 4 dpi. We then assessed the T cell responses in lungs by RBD peptide re-stimulation, and a significant increase in IFN-γ producing CD8 T cells (mean 8.45%, range 6.76-10.4%) was observed in the lungs of the mice that received PD1-RBD-DNA prime/LAIV-HK68-RBD boost regimen (Fig. 5C). Tissue-resident memory T cells (TRM) is a well-known antiviral T cell type that persists at mucosal sites (40). We found that a higher frequency of TRM cells (CD69^+^CD103^+^) (mean 30.1, range 17.6-40.9%) was identified in lung CD8 T cells in v4 mice (Fig. 5D). Moreover, a higher frequency of IFN-γ^+^ TRMs (mean 21.64%, range 17.1-26.3%) was found in v4 mice after RBD peptide re-stimulation (Fig. 5E). Correlation analysis showed negative correlations between the NAb titer and viral load (RdRp RNA/NP^+^ cells) and between the frequency of IFN-γ+ CD8 T cells and viral load in both lungs (Fig. 5F) and nasal turbinate (Fig. 5G), respectively, indicating their involvements in viral control.

**Fig. 5.**
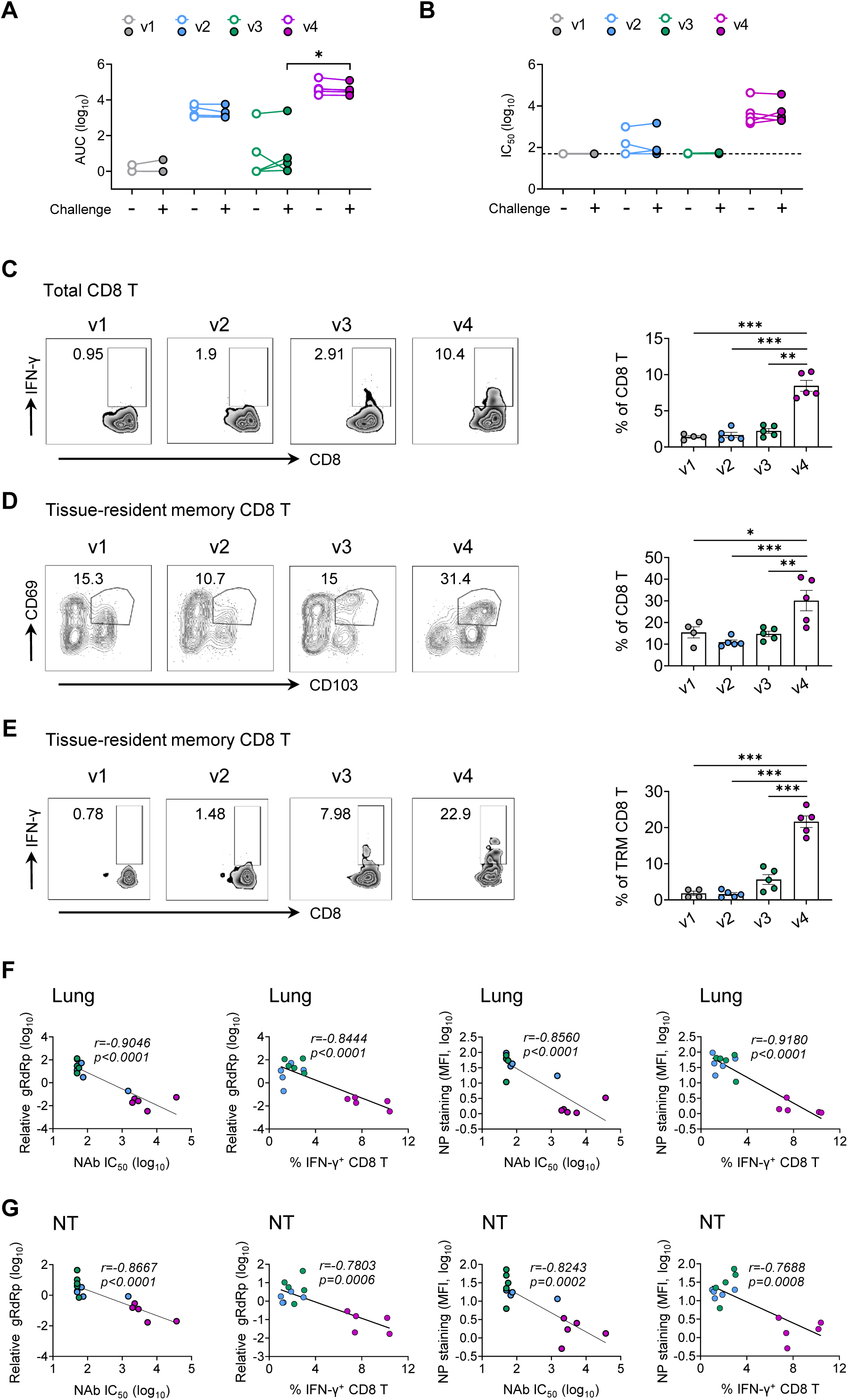
Robust NAb and antigen-specific T cells were responsible for virus control in K18-hACE2 mice. Blood samples were collected at 33 and 42 days (also 4 dpi) after the 2^nd^ vaccination for analysis. (**A, B**) The mean ± SEM changes of anti-RBD IgG AUC titer (**A**) and neutralizing IC_50_ values (**B**) were determined by anti-RBD IgG ELISA and pseudovirus assay, respectively. (**C**) Lung CD8 T cells were assayed for IFN-γ expression by flow cytometry after re-stimulated with the RBD peptide pool. Tissue-resident memory CD8 T cells (TRM) in lungs were phenotyped for expression CD69 and CD103 (**D**) and the IFN-γ producing TRM were measured by re-stimulated with the RBD peptide pool (**E**). (**F**) Viral load (relative gRdRp) in lungs correlated with NAb IC_50_ titer or percentage of IFN-γ producing CD8 T cells. NP^+^ cells (MFI) in lungs correlated with NAb IC_50_ titer or percentage of IFN-γ producing CD8 T cells. (**G**) Viral load (relative gRdRp) in NT correlated with NAb IC_50_ titer or percentage of IFN-γ producing CD8 T cells. NP^+^ cells (MFI) in NT correlated with NAb IC_50_ titer or percentage of IFN-γ producing CD8 T cells. Correlation analysis was performed by linear regression using GraphPad Prism 8.0. Each color represents a vaccination regimen. Each symbol represents an individual mouse. Statistics were generated using the 2-tailed Student’s t test. *p<0.05; **p<0.01; ***p<0.001.

### 3.8 LAIV-CA4-RBD boosted the immunogenicity of vaccines in emergency use

While we were studying the PD1-RBD-DNA/LAIV-HK68-RBD regimen, both the BioNTech mRNA vaccine and the Sinovac inactivated vaccine have been approved for emergency use in Hong Kong. Our LAIV-CA4-RBD vaccine is currently undergoing the phase I clinical trial in Hong Kong (ClinicalTrials.gov Identifier: NCT04809389). Since intramuscular administration of mRNA or inactivated vaccine induces mainly systemic IgG and some T cell immune responses without secretory IgA or tissue-resident memory T cells (41, 42), we sought to determine if LAIV-CA4-RBD could boost their mucosal immunogenicity to provide the implication for clinical use. We, therefore, immunized additional groups of BALB/c mice as v7, v9, v8, v10 with a homologous *i*.*m*. BioNTech (1/5 of clinical dose), a homologous *i*.*m*. Sinovac (1/5 of clinical dose), a heterologous *i*.*m*. BioNTech plus 10^6^ PFU LAIV-CA4-RBD, a heterologous *i*.*m*. Sinovac plus 10^6^ PFU LAIV-CA4-RBD at a 3-week interval, respectively. The selection of the 1/5 of clinical dose was based on animal studies described previously (43). The control mice in v6 group were immunized by 50 μg i.m./EP PD1-RBD DNA plus LAIV-CA4-RBD as before (Fig. 6A). At day 12 after the 2^nd^ vaccination, serum and BAL were collected and subjected to detection of RBD-specific IgG and IgA as well as neutralization activity. Interestingly, v8 and v10 mice did not show increased titers of IgG and Nab in sera (Fig. 6B, E), but had significantly increased titers of IgG (Fig. 6C), IgA (Fig. 6D) and Nab (Fig. 6F) in BAL as compared with v7 and v9. We then assessed T cell responses in lungs by RBD peptide re-stimulation. A significant increase of IFN-γ producing CD8 T cells (mean 55.58%, range 44.9-65.1% in v8 and mean 2.16%, range 1.5-2.64% in v10) was found in mice that received the heterologous boost of LAIV-HK68-RBD than mice received homologous BioNTech or Sinovac (mean 22.14%, range 8.89-30.8% in v7 and mean 0.37, range 0.1-0.66% in v9) (Fig. 6G). Furthermore, more total TRMs and RBD-specific TRMs were induced in v8 and v10 as compared with v7 and v9 (Fig. 6G-H). These results demonstrated that the intranasal LAIV-CA4-RBD boost induced stronger humoral and cellular immune responses in mucosal sites than systemic vaccinations alone.

**Fig. 6.**
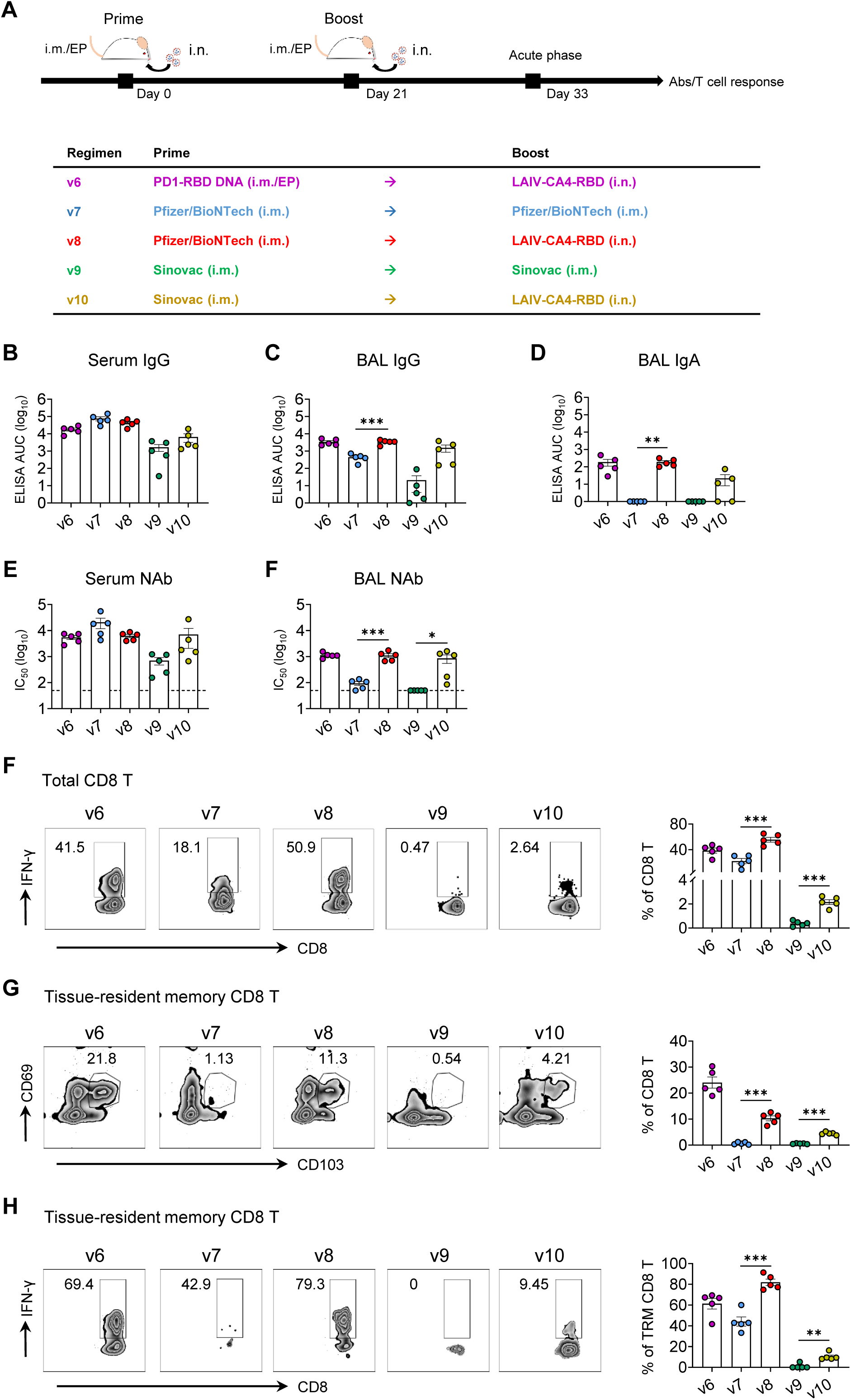
Combinations of systemic and mucosal immunization induce robust humoral and cellular immune responses in the effector site. Experimental schedule and grouping of BALB/c mice (5 mice per group). Blood (**B, E**) and bronchoalveolar lavage (BAL) (**C, D, F**) were collected and subjected to RBD-specific IgG (**B, C**) or IgA (**D**) detection and NAb IC_50_ activity (**E, F**). The area under the curve (AUC) represented the total peak area calculated from ELISA OD values. Neutralization IC_50_ values were determined against SARS-CoV-2-Spike-pseudovirus infection of 293T-huACE2 cells. (**F**) Lung CD8 T cells were assayed for IFN-γ expression by flow cytometry after re-stimulated with the RBD peptide pool. Tissue-resident memory CD8 T cells (TRM) in lungs were phenotyped for expression CD69 and CD103 (**G**) and the IFN-γ producing TRM were measured by re-stimulated with the RBD peptide pool (**H**). Each colour represents a vaccination regimen. Each symbol represents an individual mouse. Statistics were generated using the 2-tailed Student’s t test. *p<0.05; **p<0.01; ***p<0.001.

### 3.9 Vaccine-induced NAbs cross-neutralize global SARS-CoV-2 variants of concern

Recently emerged SARS-CoV-2 variants Alpha (B.1.1.7) from UK, Beta (B.1.351) from South Africa and Delta (B.1.167) from India have become challenges for passive immunotherapy and vaccine-induced protection (19, 44). For example, the Novavax vaccine is effective against the wildtype SARS-CoV-2 (95.6%) but provides reduced protection against the variants Alpha (85.6%) and Beta (60%) (45). We, therefore, compared the neutralizing activity of immune sera from PD1-RBD-DNA/BioNTech/Sinovac prime and LAIV-CA4-RBD boosted mice (Fig. 6) against pseudotyped viruses that contain the D614G, the Alpha, the Beta and the Delta variants (Fig. 7A-B) as described previously (19). Compared to the D614G viral strain, v6 sera showed slightly enhanced neutralizing activity against the Alpha variant, while the sera of other groups exhibited reduced neutralization against the Alpha, Beta and Delta variants (Fig. 7B). In line with a recent studies (19), the Beta and Delta variants were more resistant to neutralization by sera from all vaccine regimens with an average fold reduction of 1.5-1.77 (v6), 3.00-3.60 (v7), 1.98-2.03 (v8), 1.56-2.45 (v9) and 1.26-1.59 (v10) as compared to the D614G strain, respectively (Fig. 7B). Although animals in the LAIV-CA4-RBD boost regimens (v6, v8 and v10) showed the mean 1.69-fold reduction against Beta or Delta variants, their mean NAb IC_50_ titer of 1: 3633 (v6, range 1:1586-1:8250), 1:4370 (v8, range 1:2305-1:9068) and 1:3929 (v10, range 1:189-14393) remained high, respectively, which was superior or comparable to those of homologous-vaccinated mice as well as to the results of clinical vaccines against the wild type virus in murine models (6, 46, 47). Importantly, the BAL from LAIV-CA4-RBD boost groups (v6, v8 and v10) were still able to neutralize the Beta and Delta variants (v6: mean 453, range 117-762; v8: mean 280, range 93-465; v10: mean 281, range 50-760) although they showed the average fold reduction of 2.64-2.90 (v6), 1.7-4.3 (v8) and 1.26-2.28 (v10) as compared to the D614G strain, respectively (Fig. 7C-D). These results demonstrated that the systemic prime/LAIV boost regimen-induced high amounts of systemic and mucosal NAbs may confer cross-protection against the variants of concern before the tailor-made vaccines become available.

**Fig. 7.**
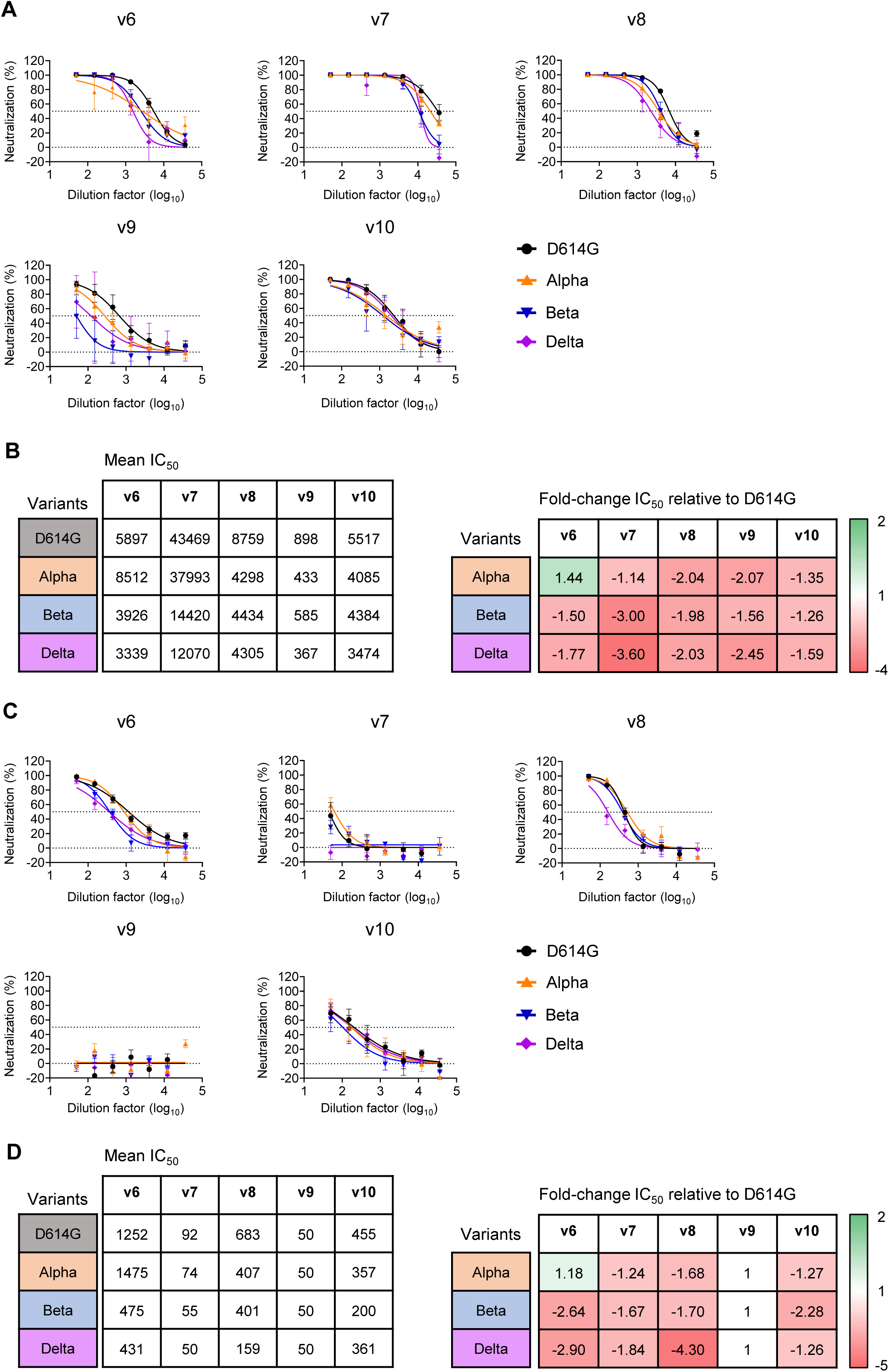
Cross-neutralization of vaccine-induced systemic and mucosal NAb against SARS-CoV-2 variants. Neutralization activities of immune sera (**A-B**) and BAL (**C-D**) elicited by various vaccine regimens (same as described in Fig. 6) were determined against 3 color-coded pseudoviruses in 293T-ACE2 cells. Data showed mean±SEM of 5 mice in each vaccine group. (**B and D**) Comparison of mean IC_50_ values against different SARS-CoV-2 variants (left) and fold-change of mean IC_50_ values in relation to the SARS-CoV-2 D614G strain (right).

## 4. Discussion

To control the ongoing COVID-19 pandemic, vaccine-induced protective immune responses should prevent SARS-CoV-2 nasal infection effectively to eliminate viral transmission between humans (17, 41). Failure of protection against SARS-CoV-2 in NT may allow asymptomatic viral spread and there have been over 10 thousand cases of vaccine breakthrough infections in the United States (45, 48). To date, little is known about the correlate of vaccine-induced mucosal protection for the prevention of intranasal SARS-CoV-2 infection in animal models (15). In this study, we demonstrated that the intranasal LAIV-based vaccination is critical for boosting the systemic PD1-RBD-DNA vaccine prime for effective prevention of SARS-CoV-2 infection in both NT and lungs by inducing high amounts of mucosal IgA/IgG and polyfunctional memory CD8 T cells. Consistent with previous findings that a heterologous prime and boost vaccine regimen may induce stronger immune responses (49, 50), our results on effective prevention of intranasal SARS-CoV-2 infection in NT would warrant the upcoming clinical development of PD1-based and LAIV-based COVID-19 vaccines (ClinicalTrials.gov Identifier: NCT04809389). Our findings may have significant implications to use LAIV-CA4-RBD for boosting the current vaccines under emergency use especially the extensively used nucleic acid-based COVID-19 vaccines (e.g. BioNTech mRNA).

The induction of mucosal immune responses is essential for preventing SARS-CoV-2 transmission effectively (39, 41). There are, however, some challenges underlying the prevention of SARS-CoV-2 in NT. First, SARS-CoV-2 exhibited a rapid burst of viral replication in NT (1), which has not been previously found with SARS-CoV (9, 51). Second, after viral entry, ciliated nasal epithelial cells with the highest expression of ACE2 and TMPRSS2 may facilitate more efficient cell-cell transmission (8), which probably makes it difficult for systemic NAb to block. Third, NAbs by systemic vaccination or passive immunization are less distributed on the surfaces of NT for the prevention of SARS-CoV-2 (17). Due to these challenges, it is critical to investigate vaccine-induced mucosal NAbs and CD8 T cells for SARS-CoV-2 prevention in NT. Thus far, most published vaccines have indicated excellent protective efficacy in the lungs of vaccinated animals without detailed analysis in NT (52-54). Several previous studies had shown no significant viral load drops in nasal swabs or nasal turbinates of vaccinated animals (6, 7, 43, 55, 56). One study found that optimal protection was achieved by only the DNA vaccine encoding the full S protein in both the upper and lower respiratory tracts against SARS-CoV-2 in rhesus macaques when vaccinated animals developed pseudovirus neutralizing IC_50_ titer less than 1:1000 (57). Serum NAb titers, as measured by both pseudovirus and live virus neutralization, served as a significant correlate of protection. Similar protection in both the upper and lower respiratory tracts was achieved by the AD26 vaccine encoding the full S protein against SARS-CoV-2 in rhesus macaques in another study (56). Both studies, however, used the macaque model that requires inoculation of 10^5^ TCID_50_ SARS-CoV-2 into each nare and intratracheal for effective infection, which is not natural and is in great contrast to the robust nature of NT infection in humans. Using a DNA vaccine encoding the full S immunogen, we recently reported that there was significant protection in the lung but not in NT against SARS-CoV-2 in Syrian hamsters even though the vaccinated animals developed pseudovirus neutralizing IC_50_ titer larger than 1:1000 (17). In this study, we consistently found that significant prevention of NT was not achieved by two systemic PD1-RBD-DNA vaccinations in both hACE2-tranduced mice and K18-hACE-2 mice.

For direct mucosal vaccination, a single-dose intranasal ChAd vaccine exhibited a significant reduction of viral loads in NT, yet the extent of infected cells in NT was not determined, and its potential for extensive human use remains unclear (39). Here, we showed the highest frequency of RBD-specific tissue-resident memory CD8 T cells in the respiratory system was found by the intranasal boost of LAIV-based vaccine after intramuscularly prime with various systemic vaccines (e.g. PD1-RBD-DNA, Pfizer/BioNTech and Sinovac). Moreover, besides RBD-specific tissue-resident memory CD8 T cells, we also observed the positive correlation between NAb IC_50_ values and BAL NAb IC_50_ values, and the negative correlation between NAb IC_50_ values and NP^+^ cells, especially in NT. It is encouraging to develop the LAIV-RBD vaccine as the boost vaccination combined with the prime of vaccines that were authorized for emergency use to achieve mucosal immunity against SARS-CoV-2. Furthermore, the role of the intranasal LAIV-RBD boost route in optimizing the extent and localization of humoral and cellular immune responses in mucosal site combined with systemic prime immunization is not clear yet and will be determined in the future study. For example, whether the intranasal LAIV-RBD boost can potently promote the antigen-specific T follicular helper cell responses as well as elicit the potent germinal center reaction for neutralizing antibody production in the mucosal site. Secondly, it is crucial to determine how the intranasal LAIV boost develops the antigen-specific tissue resident memory T cells which are not only destroy the infected cells but also recruit the innate and adaptive immune cells into the infected tissues via cytokines and chemokines (58).

In summary, our finding of prevention of NT against SARS-CoV-2 by the high amount and long-lasting mucosal NAbs and CD8 T cells induced by the heterologous PD1-RBD-DNA/LAIV-HK68-RBD regimen may have significant implication for COVID-19 pandemic control because of the existing mass-production industry for influenza vaccines. Meantime, one COVID-19 DNA vaccine has been approved for emergency use in India with promising results (59). Our PD1-RBD-DNA vaccine may prime stronger mucosal CD8 T cells responses besides benefits of inducing potent NAbs, better stability than mRNA vaccine, fast-tracked development and cost effective GMP production (60). Although intramuscular electroporation delivery has potential limitation for DNA vaccination in large populations, this approach has demonstrated the safety, tolerability and immunogenicity profile for SARS-CoV-2 DNA vaccines in clinical trials (61). Future study, however, is needed to develop non-invasive delivery techniques for DNA vaccination in humans. In addition, simultaneous or sequential co-infection by SARS-CoV-2 and A(H1N1)pdm09 caused more severe disease than infection by either virus (62), our LAIV platform may offer an opportunity of generating a human vaccine to fight both COVID-19 and influenza.

## Supporting information

Supplemental meterials

## Contributors

Z.C. and H.C. supervised two collaborative teams, conceived of and designed the study, and wrote the manuscript. R.Z., P.W., Y.-C.W and H.X. designed the experiments, analysed the data, and prepared the manuscript. R.Z., P.W., Y-.C.W., H.X., S-. Y.L., B. W-.Y.M., K. F. W., H.H., R.C-.Y.T., and S.D. conducted the immunologic and virologic assays. L.L., A.Z. and D.Z. performed the immunofluorescence staining of animal tissue section. A.J.Z. analysed tissue pathology. Q.P., N.L., B.Z., C-.Y.C and D.Y. conducted the ELISA and neutralization assay, Z.D. and K-.K.A. performed the viral RNA measurement. K.-Y.Y. provided critical comments, supports and materials. Z.C. and R.Z. have accessed and verified the underlying data reported in the manuscript.

## Acknowledgements

We thank Dr. Jincun Zhao for kindly providing the AD5-hACE2 construct, Drs. David D. Ho and Pengfei Wang for kindly providing the expression plasmids encoding for D614G, Alpha, Beta and Delta variants, and Shanghai Teresa Healthcare Sci-Tech Co., Ltd for providing the clinically certified TERESA-EPT-I Drug Delivery Device for DNA vaccination.

## Data Sharing Statement

The authors declare that the data supporting the findings of this study are available from the corresponding author upon request.

