## Supplemental meterials for "Nasal prevention of SARS-CoV-2 infection by intranasal influenza-based boost vaccination": EbioM_Supplementary Figures.pdf

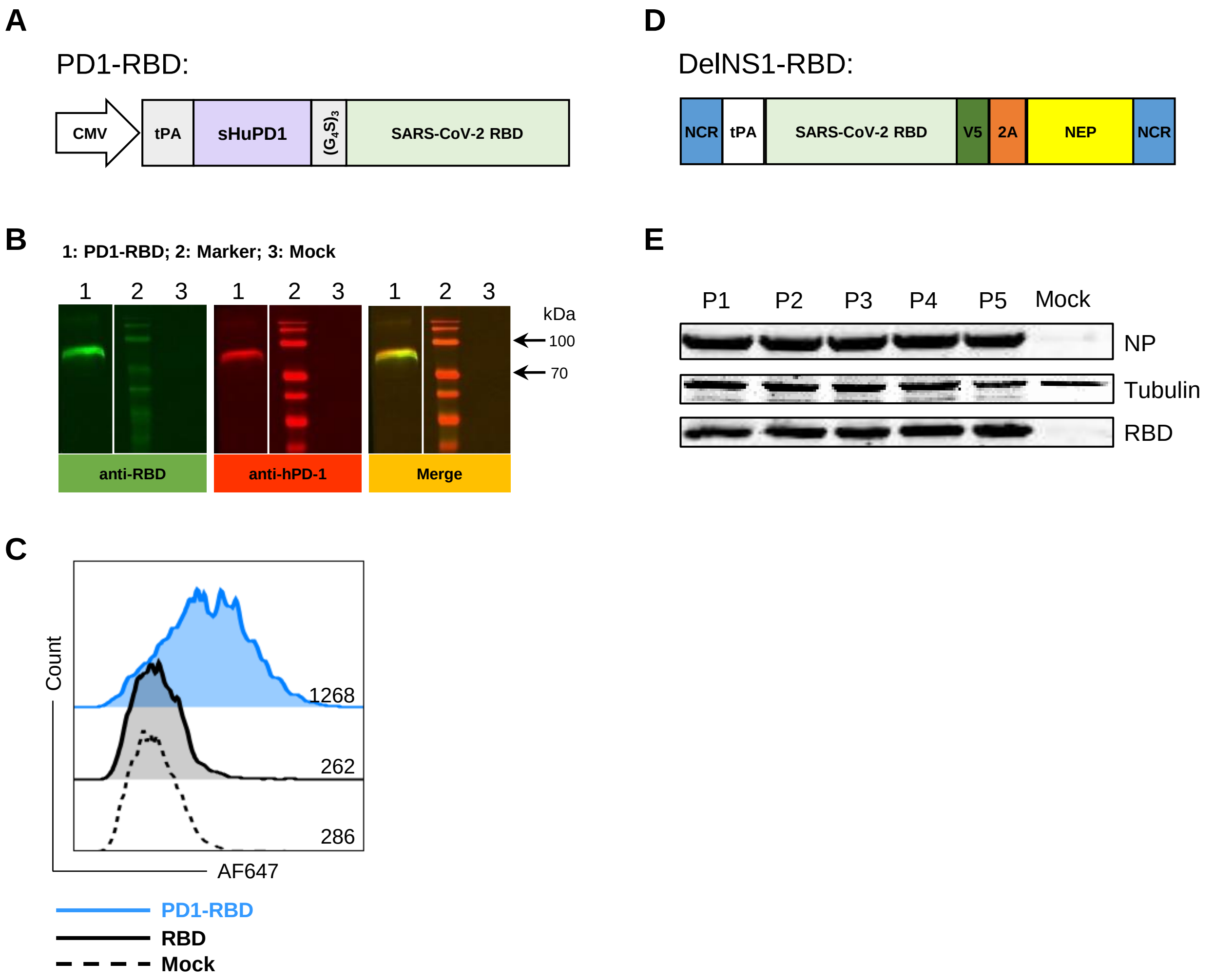


**Fig. S1. Construction and characterization of PD1-based DNA and influenza-based vaccines.** (**A**) DNA vaccine expressing SARS-CoV-2 RBD antigen fused to a human soluble PD1 domain (PD1-RBD-DNA) was constructed using the pVAX plasmid as the backbone. The protein expression was under the control of a CMV promoter and contained a human tissue plasminogen activator (tPA) secretory signal sequence to promote antigen secretion. A $(G_4S)_3$ linker sequence was placed in between the soluble PD1 domain and the antigen in PD1-RBD. The whole gene was codon optimised. (**B**) HEK 293T cells were transfected with the PD1-RBD-DNA vaccine or mock using PEI. Expression of the recombinant antigen was determined in supernatants 2 days post-transfection by Western blot analysis. The membrane was probed with rabbit anti-SARS-CoV-2 Spike antibody and mouse anti-human PD-1 antibody, respectively. The antigen-antibody complexes were detected with anti-rabbit IRDye 800CW (green) and anti-mouse IRDye 680RD (red). Lanes are identified by the legend, and the numbers in kDa indicate marker sizes (**C**) Supernatants from HEK 293T cells transfected with PD1-RBD-DNA (blue) or RBD alone (gray) were co-cultured with HEK 293T cells transiently transfected with the human PD-L1 expression vector. Supernatant from mock transfected HEK 293T cells (dashed) were used as the negative control. The binding of the recombinant antigens to PD-L1 was detected using rabbit anti-SARS-CoV-2 Spike antibodies, followed by AF647-labelled anti-rabbit secondary antibodies. Half offset histograms depict the binding of RBD. Relative geometric mean fluorescence intensity (gMFI) was shown. (**D**) SARS-CoV-2 RBD antigen was fused to an NS1 gene-deleted NS segment. The RBD domain was conjugated with a human tPA to promote antigen secretion and a V5 tag for detection (**E**) Eight pHW2000 plasmids containing the DelNS1-RBD segment and the other 7 Influenza virus genomic segments, together with an NS1 expression plasmid, were transfected into a 293T/MDCK cell mixture. Virus supernatant was collected 72 h later and designated passage 0 (P0) virus and subsequently passaged in MDCK cells 5 times (P1-P5). Cell lyses were subjected to detection of SARS-CoV-2 RBD and influenza NP.

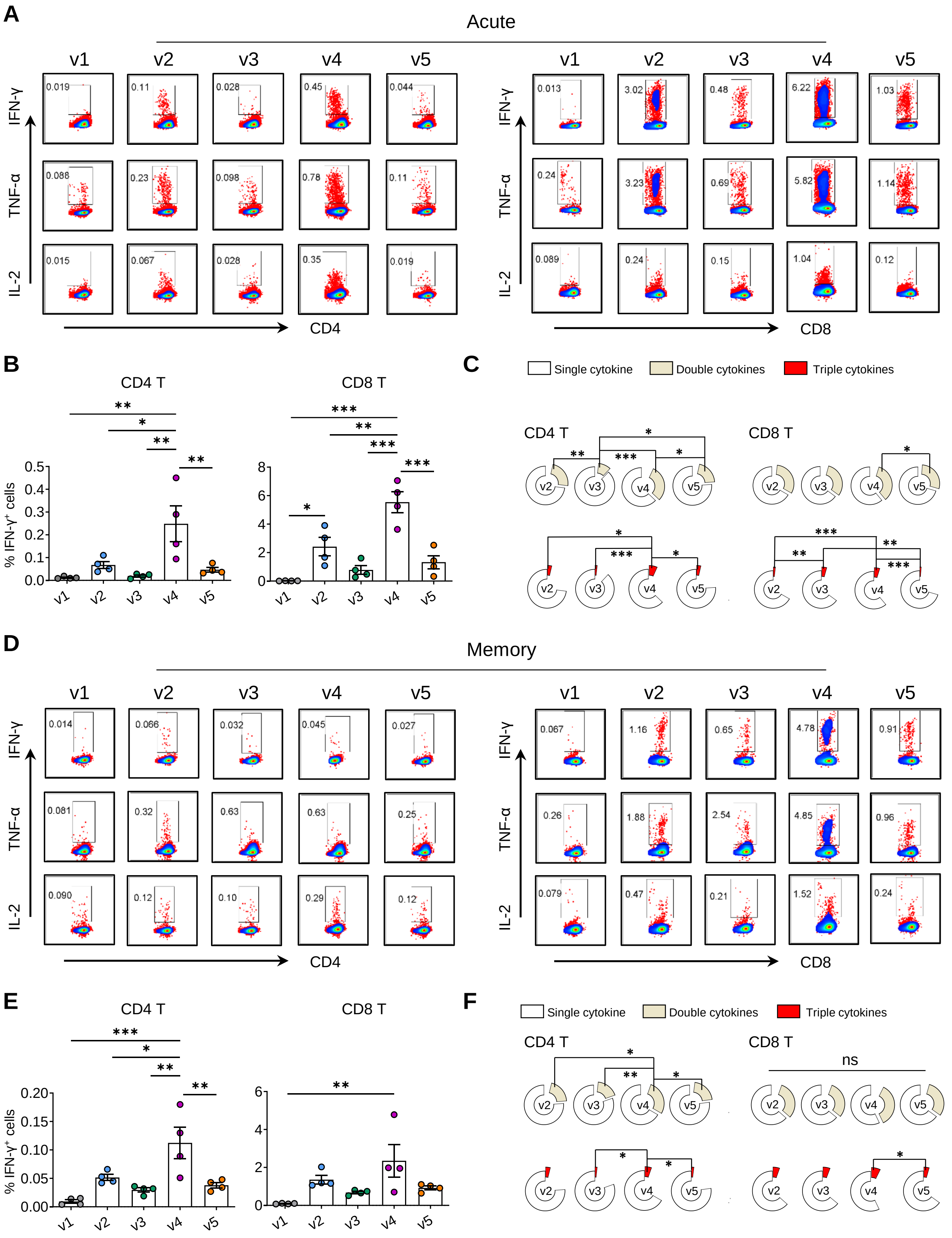


**Fig. S2. Vaccine-induced acute and memory RBD-specific T cell responses in spleen.** The vaccine immunization schedule for BALB/c mice was the same as described in Figure 1A. Splenocytes were collected and subjected to T cell response analysis at day-9 (acute) and day-48 (memory) post the 2$^{nd}$ immunization. SARS-CoV-2 RBD-specific T cell responses in the spleen were detected by *ex vivo* RBD peptide pool stimulation followed by ICS at both acute (**A-C**) and memory (**D-F**) phase. (**A, D**) Representative dot plots present the gating of IFN-γ$^+$, TNF-a$^+$ or IL-2$^+$ CD4 T (left) and CD8 T (right) against SARS-CoV-2 RBD. (**B, E**) Quantified results depict the percentage of IFN-γ$^+$ CD4 T (left) and IFN-γ$^+$ CD8 T (right). Each symbol represents an individual mouse. Error bars indicate the standard error of the mean. (**C, F**) The pie charts indicate the proportion of single or double or triple cytokines produced by CD4 T (left) and CD8 T (right). Statistics were generated using one-way ANOVA followed by Tukey's multiple comparisons test. *p<0.05; **p<0.01; ***p<0.001.

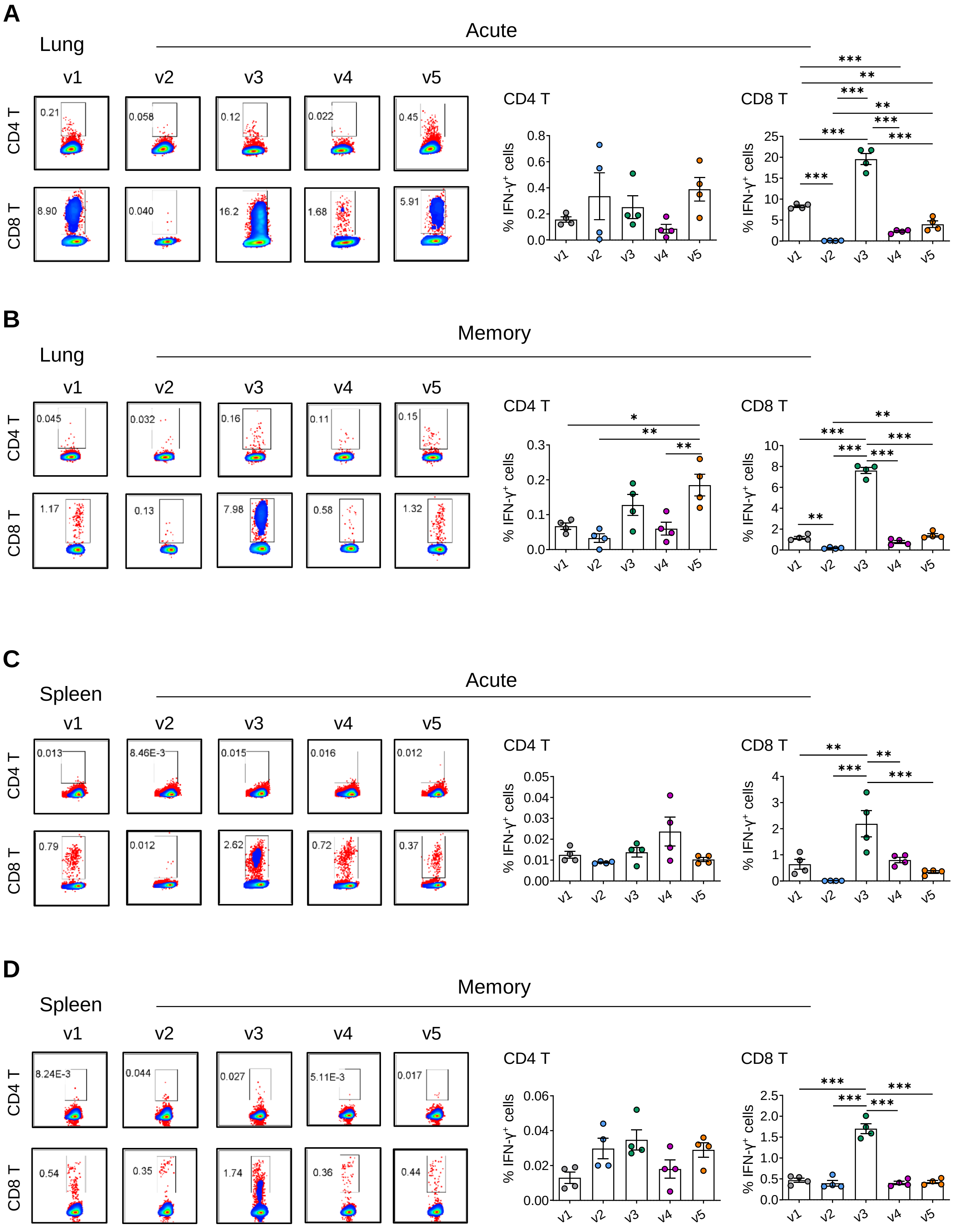


**Fig. S3. Vaccine-induced acute and memory T cell responses in lung and spleen against influenza NP.** The vaccine immunization schedule for BALB/c mice was the same as described in Figure 1A. Lung lymphocytes (**A, B**) and splenocytes (**C, D**) were collected and subjected to T cell response analysis at day-9 (acute) and day-48 (memory) post the 2$^{nd}$ immunization. Influenza NP-specific T cell responses were detected by ICS after *ex vivo* NP peptide pool stimulation at both acute (**A, C**) and memory (**B, D**) phases. Representative dot plots present the gating of IFN-γ$^{+}$ CD4 T cells (upper) and of IFN-γ$^{+}$ CD8 T cells (bottom) (**A-D,** left panel) after NP stimulation. Quantified results depict the percentage of IFN-γ+ CD4 T cells (**A-D,** middle panel) and IFN-γ+ CD8 T cells (**A-D,** right panel). Each symbol represents an individual mouse. Error bars indicate the standard error of the mean. Statistics were generated using one-way ANOVA followed by Tukey's multiple comparisons test. *$p<0.05$; **$p<0.01$; ***$p<0.001$.

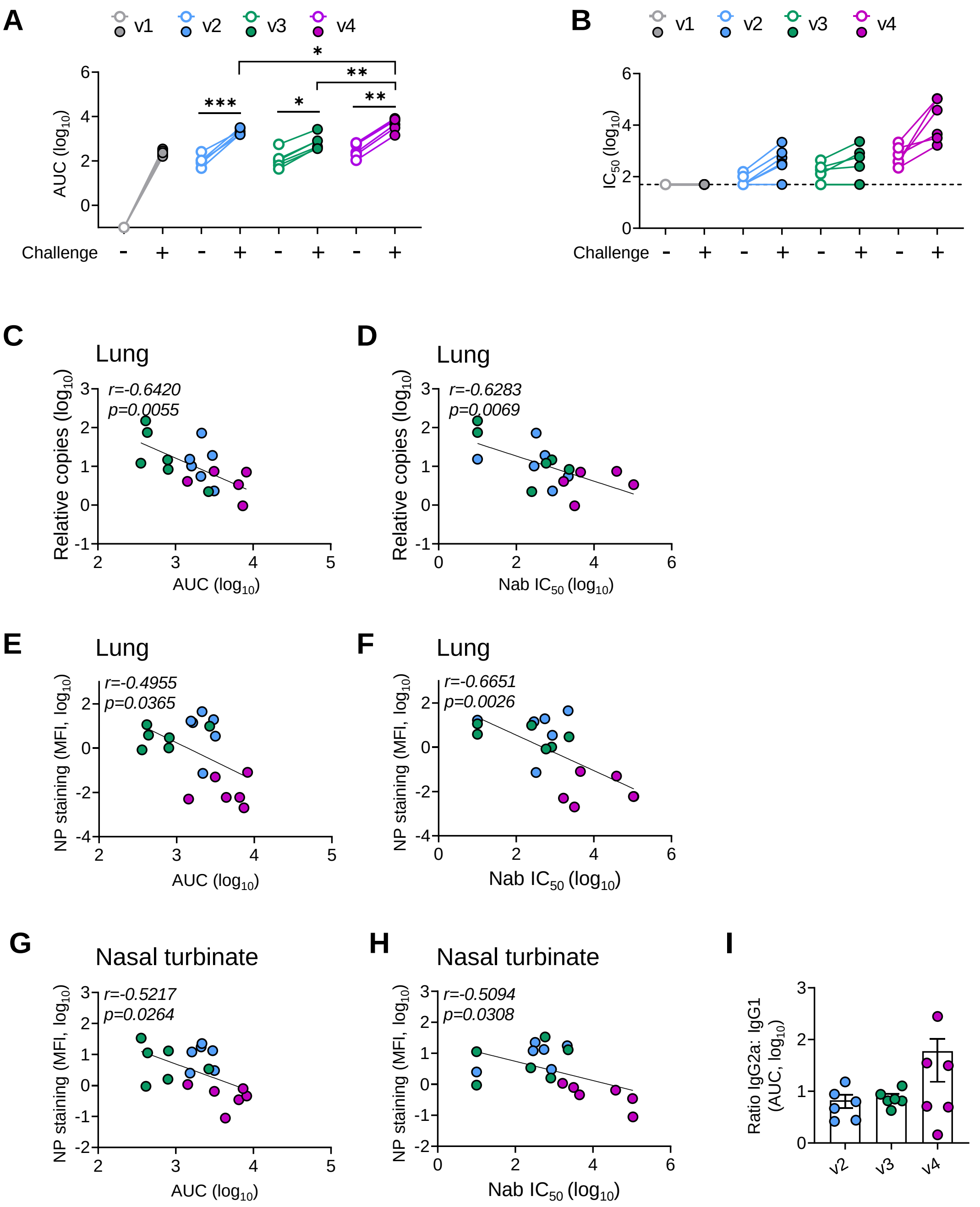


**Fig. S4. Infection-recalled NAb for correlate of protection.** Blood samples were collected at 28 and 39 days (also 4 dpi) after the 2nd vaccination for analysis. (**A, B**) The mean ± SEM changes of anti-RBD IgG AUC titer (**A**) and neutralizing $IC_{50}$ values (**B**) were determined by anti-RBD IgG ELISA and pseudovirus assay, respectively. Each color represents a vaccination regimen. Each symbol represents an individual mouse. Statistics were generated using the 2-tailed Student's t-test. *p<0.05; **p<0.01; ***p<0.001. (**C**) Viral load in lung correlated with serum IgG titer. (**D**) Viral load in lung correlated with serum NAb titer. (**E**) $NP^+$ cell in lung correlated with serum IgG titer. (**F**) $NP^+$ cell in lung correlated with serum NAb titer. Correlation analysis was performed by linear regression using GraphPad Prism 8.0. (**I**) The ratios of IgG1:IgG2a AUC titers were determined by anti-RBD IgG ELISA. Each symbol represents an individual mouse. Error bars indicate standard errors of the mean values.

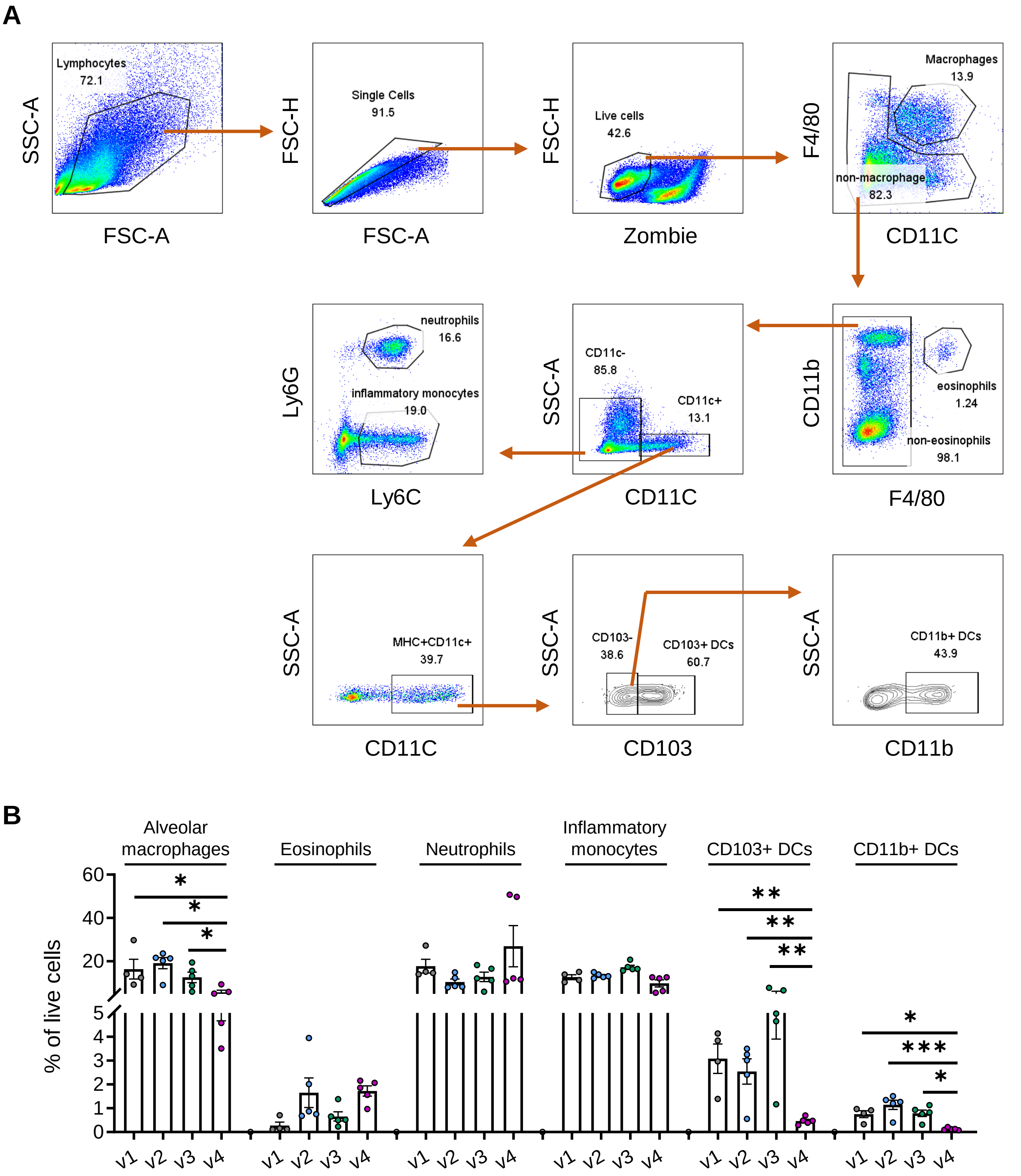


**Fig. S5. Flow cytometry analysis of immune cell composition in lung of K18-hACE2 mice at day 4 post infection.** (**A**) Flow cytometric gating strategy for lung tissue analysis. (**B**) Frequencies of different immune cells in live lung cells. Statistics were generated using the 2-tailed Student's t-test. *p<0.05; **p<0.01; ***p<0.001.
